# Pyocyanin disrupts airway regeneration to favor chronic *Pseudomonas aeruginosa* infection

**DOI:** 10.64898/2026.06.15.732350

**Authors:** Lucas A. Meirelles, Arpine Grigoryan, Laure Le Blanc, David Liaskos, Evangelia Vayena, Tania Distler, Zaïnebe Al-Mayyah, Vassily Hatzimanikatis, Alexandre Persat

## Abstract

During chronic infections, host tissues undergo repeated cycles of injury and repair while being exposed to bacterial products. Although many virulence factors trigger immune responses, whether bacterial metabolites alter tissue architecture by disrupting regenerative programs remains largely unexplored. Here, using a primary-cell-derived human lung microtissue model, we show that the redox-active metabolite pyocyanin (PYO), produced by the opportunistic pathogen *Pseudomonas aeruginosa*, disrupts airway epithelial regeneration and reshapes tissue architecture, amplifying infection. PYO exposure impairs epithelial repair, leading to defects in mucociliary clearance and promoting bacterial growth. Single-cell transcriptomics and imaging reveal that PYO exposure during basal cell differentiation alters epithelial cell-type composition. Keratin-13-rich cells promote the emergence of squamous regions with defective ciliogenesis, resulting in functional defects. After characterizing how PYO remodels the epithelium, we tested how these changes impact bacterial fitness. Using transposon-insertion sequencing and live imaging of infections, we show that these regeneration defects improve the fitness of mutants that tend to form biofilms, a common feature of chronic infections. Together, our results reveal that PYO reshapes regenerative trajectories and tissue architecture in ways that subsequently alter infection outcomes, highlighting the role of secreted metabolites as potential modulators of tissue regeneration during chronic lung infections.

## Main text

During chronic lung infections, antibiotic therapy often fails, resulting in persistent bacterial colonization near damaged and inflamed tissues (*1*, *2*). These persistent infections are hallmarks of obstructive lung diseases such as bronchiectasis, chronic obstructive pulmonary disease (COPD), and cystic fibrosis (CF). The lung environment in these patients facilitates microbial growth and exposes host cells to a diverse array of microbially secreted metabolites (*3–7*). While microbial modulation of tissue physiology via these secreted metabolites has been of intense interest in the gut (*8*, *9*), our understanding of how such interactions shape lung physiology during infections remains comparatively limited.

The opportunistic pathogen *Pseudomonas aeruginosa* causes antibiotic-recalcitrant pneumonia in individuals with obstructive lung diseases, substantially contributing to the decline in respiratory function (*10–12*). A defining feature of *P. aeruginosa* is the secretion of pyocyanin (PYO), a redox-active phenazine whose characteristic blue pigment has long served as a clinical marker of infection (*13*, *14*). PYO secretion protects *P. aeruginosa* populations from environmental stressors, such as antibiotic treatment, and likely plays a prominent role in bacterial physiology during infection (*15*, *16*).

PYO can impact host cell physiology by inducing oxidative stress, causing cell death at high concentrations (> 10 µM) (*17*), disrupting epithelial barrier integrity (*14*, *18*), impairing ciliary function (*19–21*), and stimulating mucus production (*21–24*). Mice chronically exposed to PYO show signs of lung fibrosis, goblet cell metaplasia and hyperplasia, and altered immune responses in the airway, including increased neutrophilic infiltration and damage mediated by neutrophil elastase (*18*, *21*, *25*). Yet, whether bacterial metabolites can reshape the regenerative processes that restore tissue architecture after injury, and how they may influence cell fate decisions in this context, remains largely unknown. This established knowledge positions PYO as an ideal model metabolite for addressing this question.

A hallmark of chronic infection is a localized cycle of pathogen-induced damage followed by host-driven regeneration (*26*, *27*). The lung exhibits a remarkable regenerative capacity to restore respiratory function following injury (*28*), a process governed in part by oxygen and reactive oxygen species (ROS) levels, which regulate developmental signaling pathways (*29*, *30*). In the airway epithelium, ROS levels precisely balance stem cell self-renewal and proliferation to maintain epithelial homeostasis (*31*). In lungs chronically colonized by *P. aeruginosa*, the damaged epithelia must attempt to regenerate in the presence of bacterial cells and their redox-active metabolites, including PYO (*3*, *4*, *12*, *32*). Because these metabolites alter intracellular redox states, they may intersect with redox-sensitive developmental processes required for repair (*33*, *34*).

We therefore hypothesized that chronic exposure to PYO interferes with airway regeneration. Here, we combined human airway regeneration assays, single-cell transcriptomics, high-resolution imaging, and bacterial functional genomics to determine how PYO remodels the regenerating airway epithelium and, in turn, how this environment shapes bacterial adaptation.

## Results

### PYO disrupts airway regeneration, promoting colonization

To test whether microbial redox-active metabolites affect epithelial regeneration, we examined the recovery of injured airway epithelia in the presence of PYO using human primary bronchial epithelial (HBE) cultures as a model (*28*, *35*). We physically damaged fully differentiated HBEs, then challenged them to PYO at concentrations relevant to infections (1 and 5 µM) (*32*, *33*) and monitored their injury repair (Fig. 1A). While untreated tissues regenerated into epithelia resembling their pre-injury state, PYO-exposed tissues regenerated with pronounced morphological defects (Fig. 1A). Damaged zones recovered into epithelia with variable thickness and contained cysts (Fig. 1B; Fig. S1A-B; Movie S1-2). In contrast, the morphology of undamaged HBE cells exposed to PYO remained similar to that of untreated controls (Fig. S1B), indicating that PYO specifically acts to perturb regeneration processes and spares epithelial homeostasis.

**Figure 1.**
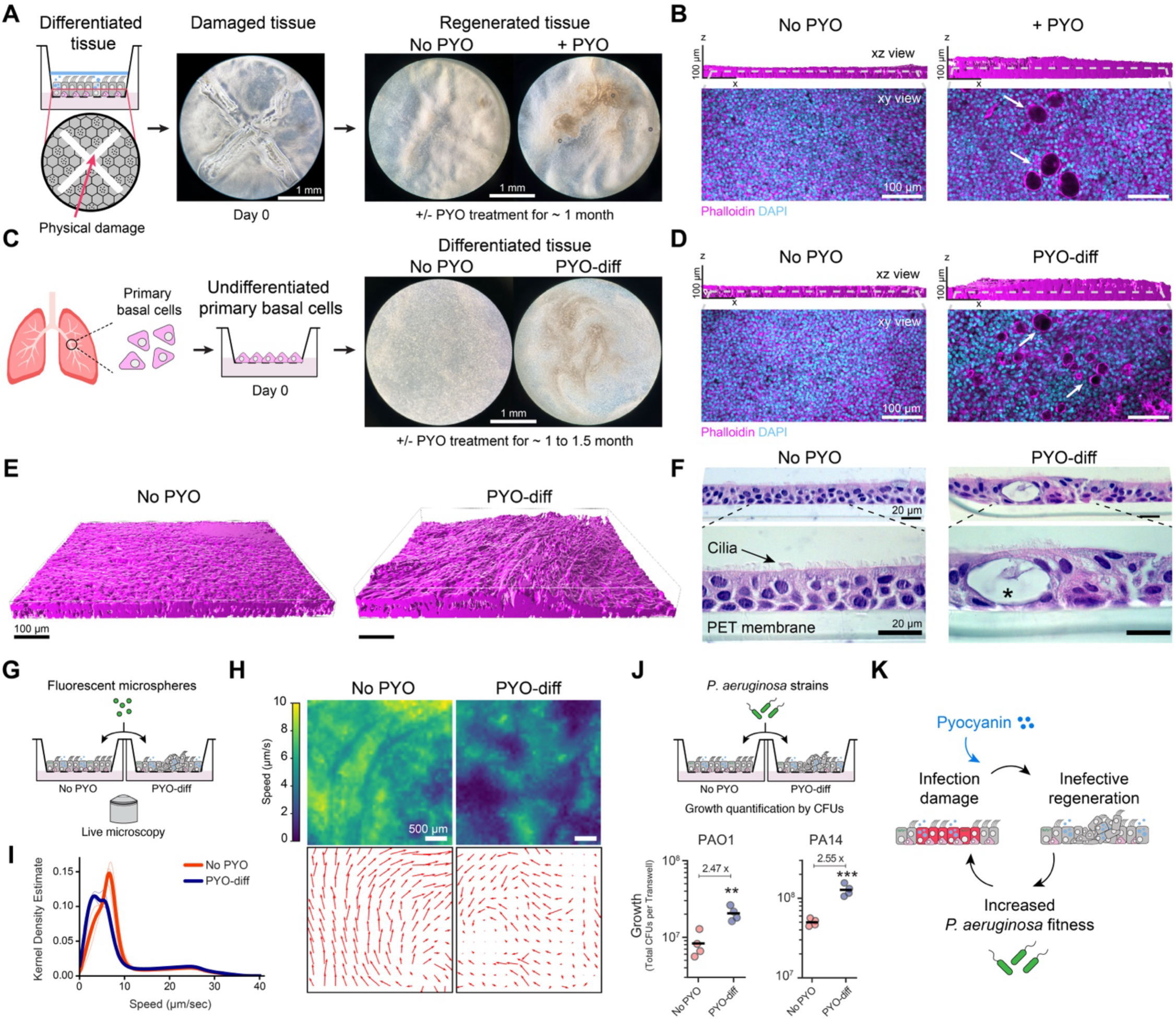
The bacterial metabolite PYO disrupts airway regeneration, promoting mucosal colonization. **A.** Human bronchial epithelial cell cultures (HBE) model of airway injury and repair. Differentiated HBE on Transwells were physically damaged and subsequently exposed to PYO (5 µM) at the basal side for 26 days. **B.** Confocal microscopy of damaged HBE after regeneration under PYO exposure (5 µM) for 29 days. Aberrant epithelial morphologies developed in the PYO-exposed sample relative to the untreated control. PYO induced tissue thickening and the emergence of cysts (white arrows). **C.** HBE differentiation in the presence of PYO exposure (PYO-diff) recapitulates aberrant epithelial phenotypes. Primary cells were seeded in Transwell inserts and differentiated with and without PYO exposure (1-5 µM) for 30-45 days. **D.** Confocal microscopy of HBE after differentiation under PYO exposure. This assay reproduced PYO-induced aberrant epithelial morphologies observed during regeneration. As in panel B, PYO induced tissue thickening and the emergence of cysts (white arrows). **E.** 3D rendering of confocal images of epithelia under PYO-diff exposure (1 µM) stained for actin. The untreated control shows uniform thickness, while the thickness is dramatically more heterogeneous in PYO-diff HBE. **F.** Histology slide (Hematoxylin and eosin - H&E - stain) of tissues differentiated with and without PYO (5 µM). The asterisk marks a cyst. **G.** Experimental design of mucociliary clearance assay by particle image velocimetry (PIV). **H**. Representative heatmaps of flow speed (top) or velocity vector fields (bottom) show that differentiation under PYO (5 µM) increases flow heterogeneity. **I.** Kernel density estimation of the distribution of mucociliary flow speed. Thin lines represent each independent biological replicate (n = 3), and thicker lines are their mean. **J.** *P. aeruginosa* growth in untreated controls vs. PYO-diff HBE. We counted *P. aeruginosa* cells (laboratory strains PAO1 and PA14) 9-10 hours post-infection. Each data point represents an independent biological replicate (n = 4); horizontal black lines mark their mean. **K.** Hypothesis of a vicious cycle of infection that may be amplified by the secretion of PYO. Experiments shown here used HBE cells from a CF donor (panel E; see Fig. S2 for additional data) and a healthy donor (all other panels). To improve the display of morphological features, the brightness and contrast in panels B, D, and E were adjusted independently for each image. Statistics in **J**, Welch unpaired t-test (two-sided) (\*\**p* < 0.01; \*\*\**p* < 0.001).

During regeneration, airway basal cells proliferate and differentiate into specialized cell types, including goblet, ciliated, and club cells, thereby restoring epithelial function (*36*, *37*). To test whether PYO interferes with this process, we differentiated primary HBE cultures, which are initially predominantly composed of basal cells, in the presence of PYO (Fig. 1C). Hereafter, we refer to this condition as PYO exposure during differentiation, abbreviated as “PYO-diff”. PYO-diff consistently induced defects resembling those observed during recovery from injury across cells from independent donors (Fig. 1C; Fig. S2A; Movies S3-4). Confocal imaging revealed aberrantly thick epithelia in regions forming cyst-like structures (Fig. 1D and 1E; Fig. S2A-B). These also correlated with irregular ciliation patterns (Fig. 1F; Fig. S2B). Perturbations in ciliary distribution and epithelial morphology may affect mucociliary clearance, a critical innate defense of the airway epithelium against microbial invaders (*2*). We therefore quantified mucociliary flow in PYO-diff HBE cultures by tracking passive fluorescent microsphere tracers in live cultures (Fig. 1G). To assess only differences in tissue structure induced by PYO during differentiation, and not immediate physiological changes such as ATP depletion and the resulting reduction in cilia beating frequency (*21*), the molecule was removed ∼24 hours before the experiment. Particle image velocimetry analysis shows that untreated cultures generate a coherent mucociliary flow with uniform speed, indicating robust clearance. By contrast, flows generated by PYO-diff HBE cultures exhibit lower coherence and greater speed variations, resulting in flow patterns with stagnant patches (Fig. 1G-I, Fig. S3; Movie S5-6).

In individuals with chronic airway disease, disruption of mucociliary clearance promotes mucus accumulation, creating microenvironments that favor pathogen growth (*2*). To test whether *P. aeruginosa* can benefit from PYO-induced clearance defects, we measured the growth of *P. aeruginosa* laboratory strains PAO1 and PA14 in PYO-diff HBE cultures. Both strains grew significantly better in PYO-diff tissues relative to untreated controls (Fig. 1J). Thus, by secreting PYO during chronic infections, *P. aeruginosa* could disrupt epithelial regeneration and impair airway mucociliary clearance, thereby benefiting colonization. We hypothesized that this process may establish a vicious cycle in which enhanced bacterial fitness leads to a higher bacterial burden, resulting in locally elevated PYO levels and increased tissue damage (Fig. 1K) (*26*, *27*). To elucidate the molecular mechanisms driving this self-reinforcing process, creating these favorable microenvironments, we next investigated how PYO influences epithelial differentiation.

### PYO modulates cell-type composition, leading to the emergence of a subcluster of ciliated cells

Epithelial overgrowth and heterogeneous mucociliary flow patterns indicate that PYO perturbs epithelial differentiation. We thus reasoned that PYO could impact the composition of airway cells, which we could type by single-cell RNA-sequencing (scRNA-seq) in both untreated (“NC”) and PYO-diff HBE cultures (1 µM, “PYO1”; or 5 µM, “PYO5”) (Fig. 2A). To distinguish developmental effects of chronic PYO exposure from acute physiological responses, we also analyzed transcriptomes of fully differentiated epithelial cultures exposed to 1 µM PYO for shorter durations (“PYO-shock”). Single-cell transcriptomes showed that all the tissues contained canonical airway epithelial cell types, including basal, club, goblet, ionocyte, deuterosomal, and ciliated cells, as well as KRT13^+^ cells resembling hillock cells (Fig. 2B; Fig. S4A-F) (*36*, *38*, *39*).

**Figure 2.**
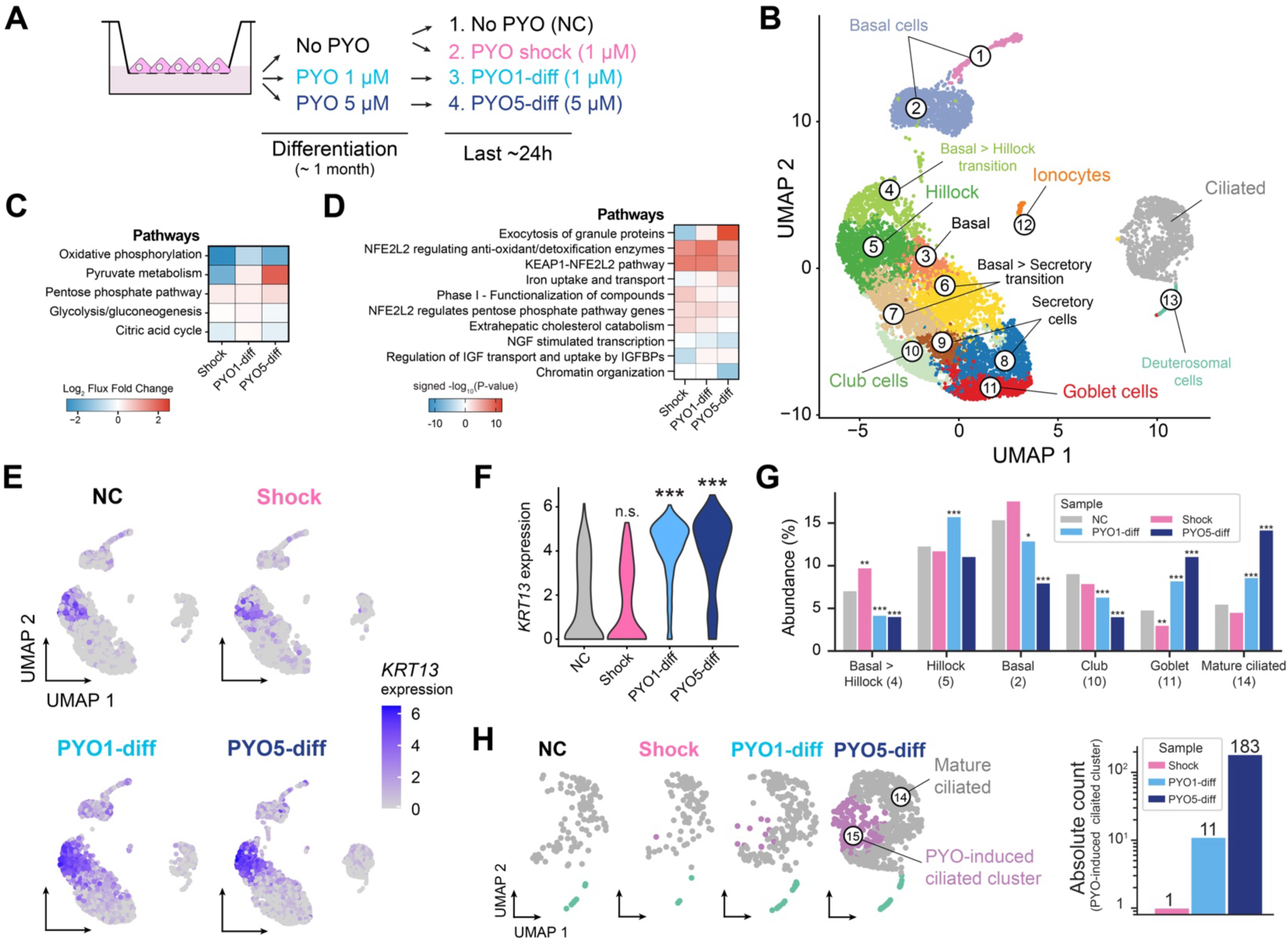
PYO modulates cell-type composition, leading to the emergence of a subcluster of ciliated cells. **A.** Studying how PYO affects tissue development with single-cell RNA sequencing (scRNA-seq). An experimental design was used to assess the effects of PYO on tissue differentiation and cell type composition. Four different treatments were prepared: 1. Cells never encountered PYO throughout the differentiation process; 2. Cells were exposed to a shock of PYO (1 µM) only in the last ∼24 h of the experiment, after the tissue was already differentiated; 3-4. Cells were chronically exposed to PYO throughout the entire differentiation process at lower or higher doses (1 or 5 µM; PYO1-diff or PYO5-diff, respectively). **B.** Uniform Manifold Approximation and Projection (UMAP) of 10163 cells assessed with scRNA-seq. Different colors represent distinct clusters, which were then annotated post hoc. **C.** Heatmap depicting the average metabolic flux deregulation (log₂ fold-change) of reactions within central carbon and energy metabolic subsystems across three treatments relative to the negative control, as inferred from the generated metabolic model. See Table S1-S2 for additional information. **D.** Enrichment of the top 10 signaling pathways at the population level for each of the three treatments versus the negative control. Heatmap displays Benjamini-Hochberg adjusted-log_10_ (*p*-values) calculated via upper-tailed hypergeometric test, signed to indicate enrichment of upregulated (red) or downregulated (blue) genes. See Table S3-S4 for additional information. **E-F.** *KRT13* expression levels across the four conditions studied are shown in a UMAP of all cells (E, with high expression in cells assigned to the basal hillock and hillock clusters), or in violin plots of only cells from clusters 4 and 5 (F). For data on the two clusters separately, see Fig. S4G. **g.** Changes in cell-type composition across the different treatments. **h.** UMAP highlight of ciliated (FOXJ1^+^) cells split by treatment (left) and quantification of the number of cells within the subcluster induced by PYO, colored in purple (right). The scRNA-seq experiment used HBE cells from a healthy donor – see Methods for details. Statistics in **F**, Wilcoxon Rank Sum test after Benjamini-Hochberg correction for multiple comparisons, with asterisks showing significant differences relative to NC (\*\*\**p* < 0.001; n.s., *p* > 0.05); in **G**, test of equal or given proportions, with asterisks showing significant differences relative to NC (\**p* < 0.05; \*\**p* < 0.01; \*\*\**p* < 0.001; n.s., p > 0.05).

As a redox-active molecule, PYO has the potential to impact cellular metabolism and signaling (*33*, *34*). To identify acute redox responses and distinguish them from developmental processes, we compared metabolic pathways enriched under PYO-shock and PYO-diff conditions. Metabolic flux analysis revealed that PYO-diff shifted central carbon metabolism, with increased flux through pyruvate metabolism compared with untreated controls. Pyruvate metabolism was, in contrast, repressed under PYO-shock conditions (Fig. 2C; Fig. S5A-B). Analysis of signaling pathways showed that both PYO-shock and PYO-diff exposures activated canonical xenobiotic detoxification programs, including Nrf2-mediated oxidative stress responses (Fig. 2D; Fig. S5C-D). PYO-diff, however, induced pathways in a cell-type-specific manner that were not activated in PYO-shock (Fig. S5D). These included pathways associated with iron uptake and transport, most significant in secretory, goblet, and ciliated cells; cell adhesion, most significant in basal and basal-to-hillock cells; and exocytosis in all but basal cells (Fig. S5D). Moreover, PYO-diff uniquely activated β-catenin-independent Wnt signaling in secretory cells, retinoic acid signaling in most cell types, and multiple interleukin signaling pathways, particularly in basal-to-hillock and hillock cells (Fig. S5D). β-catenin-independent Wnt and retinoic acid signaling are tightly involved in tissue differentiation (*40*, *41*). Together, these results indicate that PYO exposure during epithelial differentiation modulates metabolic and signaling programs distinctly from those triggered by acute exposure. The nature of signaling cascades in PYO-diff tissues further suggests that differentiation pathways have been altered, providing a basis for downstream changes in the cell-type composition of these tissues.

Alterations in retinoic acid metabolism are characteristic of basal hillock cells, which display upregulated retinoic acid catabolism and increased sensitivity to its signaling inhibition (*39*). We therefore examined how PYO-diff affected cells exhibiting hillock-related transcriptional characteristics in our dataset. Two of the detected clusters (4 and 5) expressed the hillock marker KRT13 (Fig. 2E-F; Fig. S4G). Although we did not observe dramatic changes in the proportion of these cells across treatments (Fig. 2G; Fig. S4F), PYO-diff HBE cultures showed a greater fold change in *KRT13* expression (Fig. 2E-F; Fig. S4G). In addition, these cultures had fewer basal and club cells, but a higher proportion of goblet and ciliated cells than in untreated and PYO-shock controls (Fig. 2G; Fig. S4F). Within the increased ciliated cells, we noted a subpopulation that appeared only when cells were exposed to PYO during differentiation, with a unique subcluster emerging in PYO-diff HBE cultures (Fig. 2H). Together, our results demonstrate that PYO exposure during airway epithelial regeneration perturbs basal cell differentiation, reshaping cell-type composition and inducing abnormal tissue morphology and ciliation.

### PYO disorganizes ciliation patterns

A subcluster of ciliated cells emerged in PYO-diff HBE cultures. Despite the expression of the transcription factor FOXJ1 as a ciliated cell marker, this subcluster is characterized by reduced expression of genes involved in cilia organization, assembly, and movement (Fig. 3A-B; Fig. S4H). To test whether this transcriptional signature caused defective ciliogenesis, we co-stained tissues for FOXJ1 and the cilia marker α-tubulin. Immunofluorescence shows that in PYO-diff tissues, many FOXJ1⁺ cells lack cilia (Fig. 3C-D). Cells that display cilia were sparsely distributed, producing large unciliated patches contrasting with the homogeneous ciliation observed in untreated controls (Fig. 3C). Thus, despite FOXJ1 expression, these cells did not exhibit the archetypal ciliated phenotype, leading us to re-evaluate assignment.

**Figure 3.**
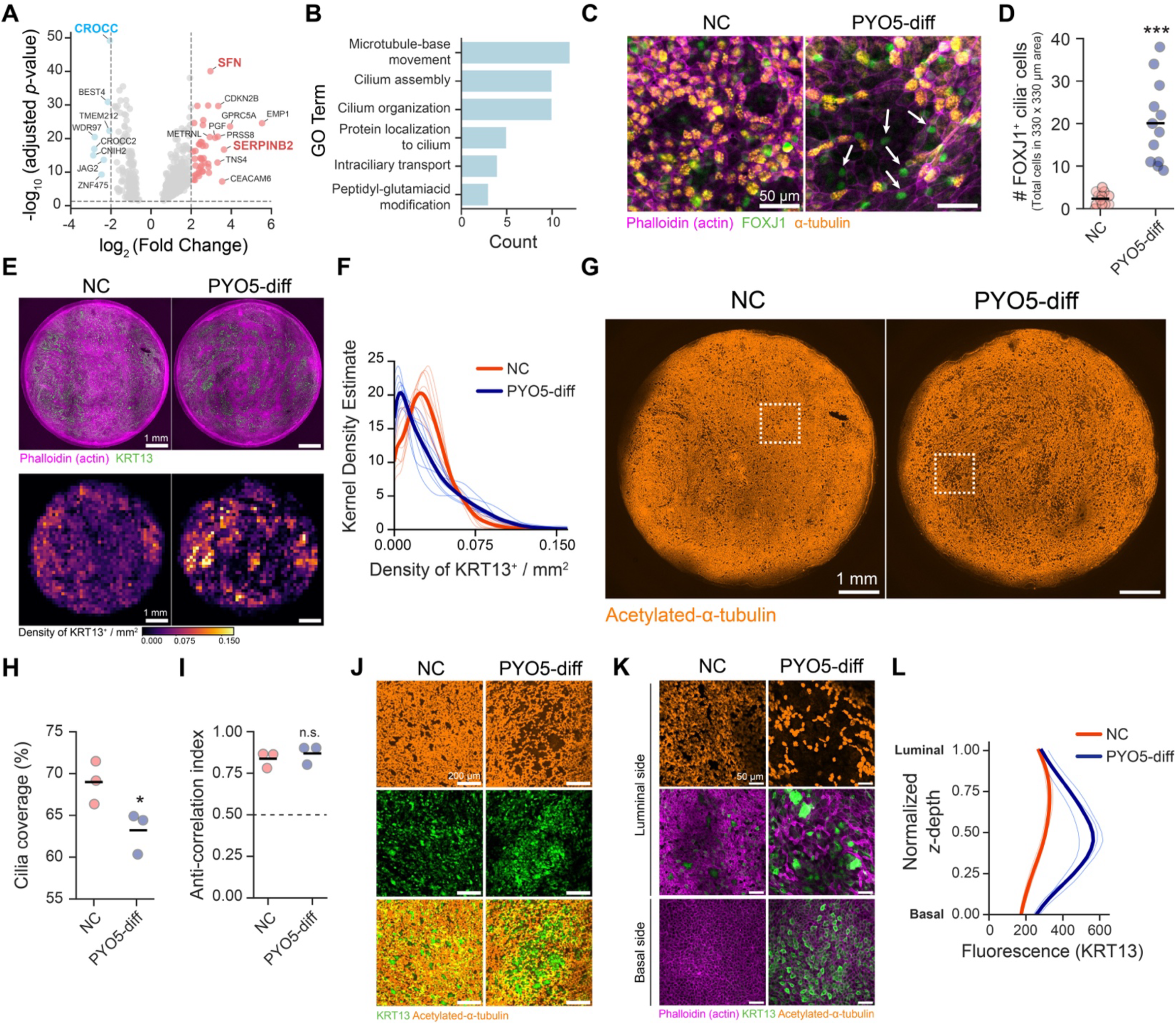
PYO disorganizes ciliation patterns. **A.** Differential gene expression between PYO-induced ciliated cells and mature ciliated cells (clusters 15 vs. 14 in Fig. 2H, respectively). Only cells from PYO5-diff treatment were used. Volcano plot with genes up-and down-regulated. CROCC (cilia assembly (*63*)), SFN (squamous marker (*42*)), and SERPINB2 (one of the hillock cell markers (*39*)) are highlighted. **B.** Enrichment analysis of gene ontology (GO) terms in downregulated genes. **C.** Representative IF images showing the luminal side of airway epithelium with staining for the ciliated-cell marker FOXJ1 and cilia. Control and areas displaying altered ciliation within PYO5-diff treatment are shown, with arrows pointing to examples of FOXJ1^+^ cells lacking cilia. **D.** Quantification of the number of FOXJ1^+^ cells not displaying cilia in our images. Each data point represents one field of view (n = 12) selected among three biological replicates; horizontal black lines mark their mean. **E.** Representative images showing the spatial distribution of KRT13^+^ expression across airway epithelia differentiated with and without exposure to PYO. Top images display IF of the entire tissue, while bottom images display spatial quantification of KRT13 signal through heatmaps. **F.** Kernel density estimation of the distribution of the KRT13 signal from heatmaps. Thin lines represent each independent biological replicate (n = 8 for NC; n = 9 for PYO5-diff), and thicker lines are their mean. Note shift towards very low or very high levels in PYO-treated samples, characteristic of patchy expression compared to more evenly distributed controls. **G.** IF images showing the spatial distribution of cilia across airway epithelia differentiated with and without exposure to PYO (same samples as in E). Highlighted areas in squares are shown in J. **H.** Quantification of cilia coverage in controls and PYO-treated samples. Each data point represents an independent biological replicate (n = 3); horizontal black lines mark their mean. **I.** Anti-correlation between KRT13 and acetylated-α-tubulin in the same samples. Values above 0.5 indicate minimal overlap, with areas displaying either one of the markers. Each data point represents an independent biological replicate (n = 3); horizontal black lines mark their mean. **J.** Zoomed in areas (highlighted in squares) from panel G, together with the KRT13 signal. Note the squamous area in PYO5-diff treatment displaying dispersed ciliation, which colocalizes with patches of tissue displaying high KRT13. **K.** Representative images with higher magnification of KRT13 expression in the basal and luminal sides of tissue in the two different conditions. Control and areas displaying altered ciliation within PYO5-diff treatment are shown. Acetylated-α-tubulin signal from the luminal side is also shown. **L.** Quantification of the KRT13 signal throughout the z-depth of the tissue in the control and areas with altered ciliation within PYO5-diff treatment. Thin lines represent each independent biological replicate (n = 3), and thicker lines are their mean. All experiments shown used HBE cells from a healthy donor – see methods for details. Statistics in **D**, **H**, and **I**, Welch unpaired t-test (two-sided) (\**p* < 0.05; \*\*\**p* < 0.001; n.s., *p* > 0.05).

In addition to lacking cilia, FOXJ1⁺ cells exhibited features associated with squamous tissues. First, they were predominantly found in the luminal side of clustered patches containing basal and luminal elongated and flat cells with squamous morphology (*39*) (Fig. S6A-B). Their high expression of SFN further supports their squamous properties (*42*) (Fig. 3A; Fig. S4H). Second, relative to mature ciliated cells, FOXJ1^+^ cells displaying defective ciliogenesis strongly upregulate SERPINB2, a marker previously described for squamous luminal hillock cells (*39*) (Fig. 3A, Fig. S4H). Although these FOXJ1⁺ cells did not express *KRT13* themselves (Fig. 2E), this cytokeratin is a marker frequently found in squamous tissues (*39*, *43*, *44*). Given the notable increase of KRT13 in PYO-diff samples (Fig. 2E-F; Fig. S4G), the association of this marker with squamous tissues (*39*, *43*, *44*), and the observed disruption of ciliation in the FOXJ1⁺ cell population in these samples (Fig. 3A-D), we suspected a connection among these three characteristics. Specifically, we propose that PYO-diff promotes the expansion of basal squamous KRT13^+^ cells, resulting in localized patches lacking functional ciliated cells.

To test this hypothesis, we measured the spatial distribution of KRT13 and acetylated α-tubulin within the same tissues. Mapping KRT13 expression revealed pronounced spatial clustering in PYO-treated tissues (Fig. 3E). Density distributions show that the KRT13 signal is more heterogeneous, with an increased proportion of low and high expression relative to a more uniform untreated control (Fig. 3E-F). The ciliation of PYO-diff tissues was also, on average, lower than that of untreated controls (Fig. 3G-H). KRT13 and acetylated α-tubulin signals were largely segregated, indicating that cells typically expressed one marker or the other (Fig. 3I). Supporting our hypothesis, the regions depleted from cilia were enriched in KRT13^+^ cells (Fig. 3J-K), and within these regions, KRT13 expression extended from basal to luminal layers (Fig. 3K-L; Fig. S6B; Movie S7), with elongated flat cells exhibiting squamous morphology across these patches (Fig. S6B; Movie S7). By contrast, untreated tissues exhibited homogeneous ciliation and lower KRT13 expression restricted to uniformly distributed luminal cells (Fig. 3K-L; Movie S8).

### PYO causes heterogeneous patterns of proliferation in the epithelium

The thickening of certain regions in the epithelium containing cysts is another key characteristic of PYO-diff tissues (Fig. 1D-F; Fig. S2). We asked whether increased proliferation could contribute to the emergence of the observed epithelial thickening in PYO-diff treatments. Specifically, we tested whether PYO-diff tissues indeed exhibited altered proliferation patterns across tissues, which may have implications for tissue structure. Untreated tissues exhibited a sparse, uniform distribution of the proliferation marker Ki-67 (Fig. 4A-B; Fig. S6C). In contrast, PYO-diff tissues displayed a heterogeneous pattern, with hyperproliferative patches that mirrored the spatial pattern observed for KRT13 (Fig. 4A-B; Fig. 3E-F). The increased proliferation was localized in large clusters of Ki-67^+^ cells within thin epithelial regions adjacent to thick cystic areas (Fig. 4C; Movie S9). Based on their cell morphology and location, these highly proliferative regions correspond to thinner squamous tissue patches that exhibit high KRT13^+^ expression and reduced ciliation, often adjacent to thicker cyst-containing areas. (Fig. S6D). Together, these results suggest that PYO-mediated remodeling promotes the development of localized patches marked by disrupted ciliation, higher KRT13 expression, and increased proliferation, resulting in a heterogeneous epithelial structure with a potentially altered tissue microenvironment.

**Figure 4.**
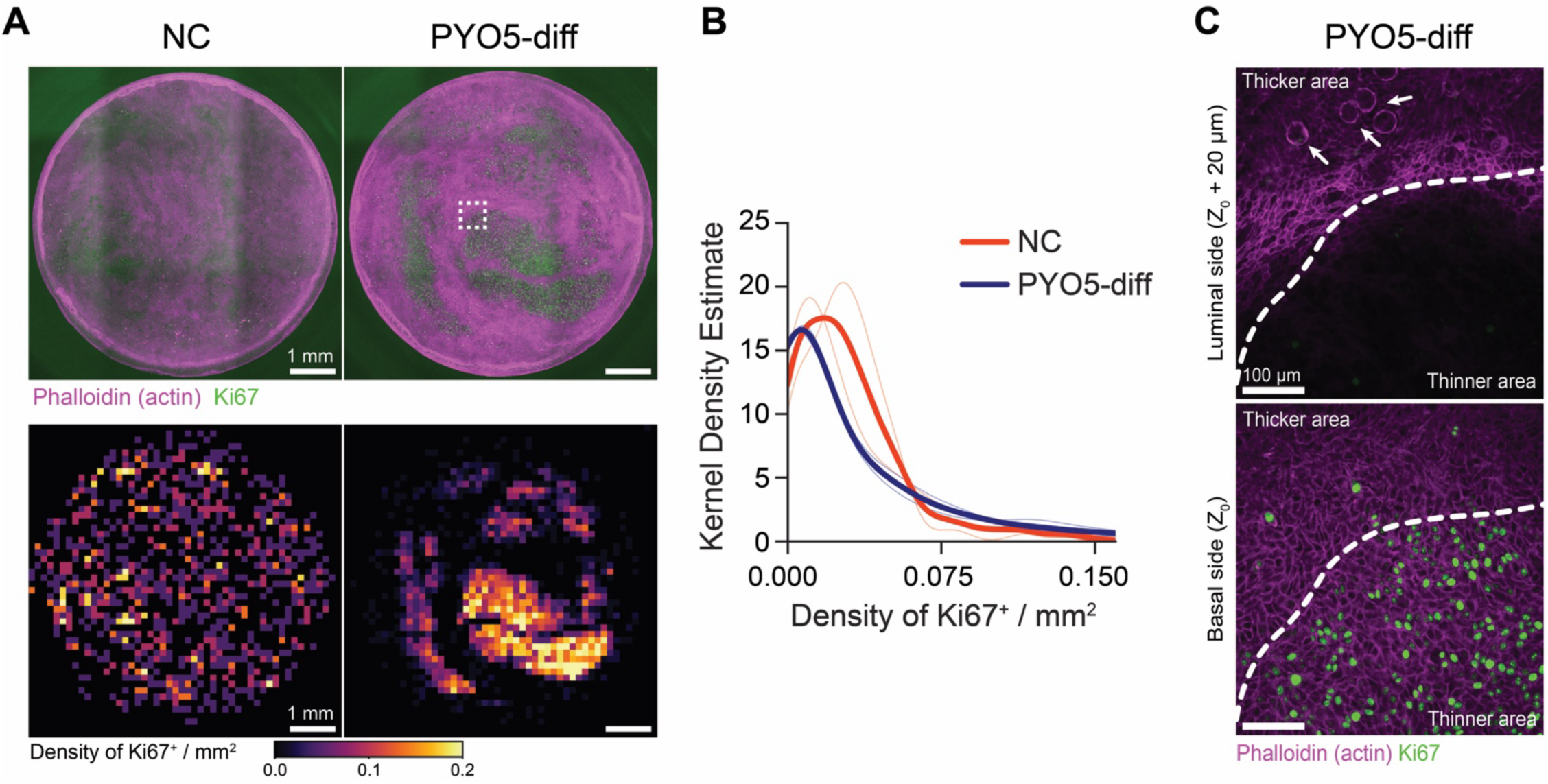
PYO causes heterogeneous patterns of proliferation in the epithelium. **A.** Representative images showing the spatial distribution of the expression of the proliferation marker Ki67^+^ across airway epithelia differentiated with and without chronic exposure to PYO. The top images IF of the entire tissue, while the bottom images display quantification of the Ki67 signal through heatmaps. The square highlights the approximate area shown at higher magnification in panel C. **B.** Kernel density estimation of the distribution of the KRT13 signal from heatmaps. Thin lines represent each independent biological replicate (n = 2), and thicker lines are their mean. As in Fig. 3F, note the shift towards very low or very high levels in PYO-treated samples, characteristic of patchy expression compared to more evenly distributed controls. **C.** Higher magnification of the highlighted square in panel A, showing the location of a large patch of proliferative cells (Ki67^+^) adjacent to thicker tissue (see Movie S9 for the full z-stack). Arrows point to cysts found in the thicker part of the tissue. All experiments shown used HBE cells from a healthy donor – see methods for details.

### Bacterial adaptation to PYO-induced developmental perturbations

Since *P. aeruginosa* grows more in PYO-diff conditions (Fig. 1J), we next investigated the mechanisms of improved bacterial colonization of remodeled tissues. To achieve this, we used transposon insertion sequencing (Tn-seq) to identify the genetic requirements for growth across the *P. aeruginosa* genome. We grew a transposon mutant library of *P. aeruginosa* PAO1 at the mucosal surface of PYO-diff HBE cultures (1 µM, “PYO1-diff”; or 5 µM, “PYO5-diff”) (Fig. 5A). As controls, we included untreated tissue (“NC”) as well as rich medium (LB) and cell-culture medium (P-ALI) to control for potential leaks (i.e., media leaking from the basal side into the air-liquid interface of the mucosal surface) in HBE cultures during infection (Fig. 5A). Tn-seq identified a core set of 156 genes essential for growth on airway tissues compared to LB (Table S5). Among these, genes related to amino acid and nucleotide biosynthesis and iron acquisition were consistently required at the mucosal surface. This supports our earlier findings of significant metabolic adaptation when shifting to growth at the mucosal surface (*45*). In addition, the fitness profile of the P-ALI control differed markedly from that of all tissue samples, suggesting minimal diffusion of basal medium to the mucosal surface (Fig. 5B). This confirms that differences between PYO-treated and untreated tissues are not due to *P. aeruginosa* feeding on basal medium.

**Figure 5.**
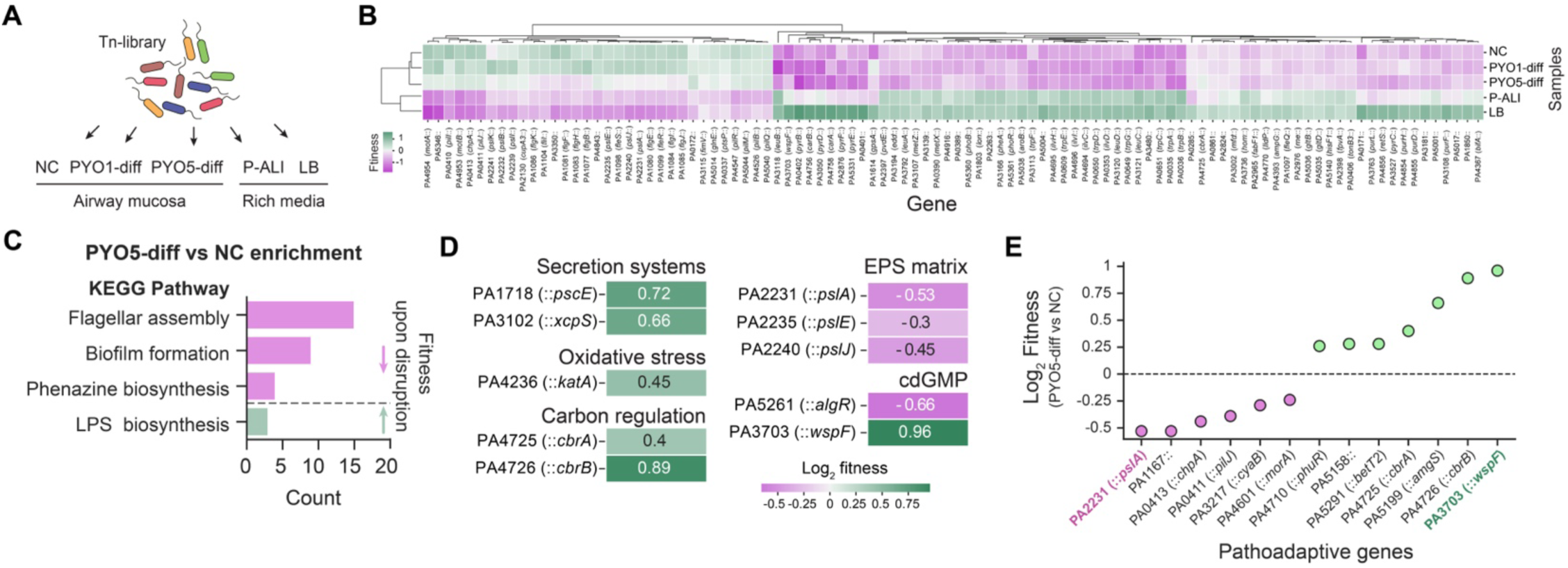
PYO-induced developmental perturbations change the fitness landscape. **A.** We use Tn-seq to identify the fitness landscape of *P. aeruginosa* during growth in HBE exposed to PYO during differentiation (Negative untreated control – NC, no PYO added; PYO1-diff, 1 µM added; PYO5-diff, 5 µM added). PYO was washed away before infection with the transposon library. Two rich media were used as additional controls (Luria-Bertani – LB, used to prepare and grow the library; PneumaCult-ALI medium – P-ALI, used to differentiate and feed the tissue from the basal side). Two independent replicates were performed for each sample. For full experimental design, see the methods section. **B.** Fitness of top-100 genes identified as significant with one-way ANOVA (adj. *p*-value < 0.05, performed with TRANSIT; see Table S8 for full list) across the five treatments, clustered by similarities. Fitness values in this plot are log-fold-changes of counts in each condition relative to the mean read counts across all conditions (see Table S8). **C.** Enrichment analysis of the Tn-seq results comparing PYO5-diff to NC tissues (KEGG pathway; see Table S7 for details). Disruption of processes in magenta decreased fitness, while disruption of processes in green increased fitness. An additional term associated with decreased fitness was “Two-component system”, but we did not include it in the figure due to its lack of specificity. **D.** Fitness effects of transposon insertions in representative genes related to chronic behavior identified by Tn-seq (PYO5-diff compared to the NC control; see Table S6 for full list). **E.** Fitness effects of transposon insertions in genes recognized as pathoadaptive that made the significance cutoff (adj. *p*-value < 0.1) (*46*). Notice the opposite effects of disruptions of *pslA* and *wspF*.

To test whether PYO-diff resulted in distinct selective pressures on *P. aeruginosa*, we compared the fitness of mutants under perturbed and normal tissue conditions. We observed mild fitness changes in PYO1-diff tissues (a total of 58 genes passed the threshold for significant fitness changes, 18 with reduced and 40 with increased fitness, adj. *p* < 0.1) (Table S6). PYO5-diff tissues displayed more pronounced shifts, with 225 significant mutants (83 with reduced and 142 with increased fitness, adj. *p* < 0.1) (Table S6). Enrichment analysis in the PYO5-diff results identified pathways associated with motility, biofilm formation, phenazine production, and LPS biosynthesis (Fig. 5C, Table S7). PYO5-diff tissues also favored mutations involved in secretion systems and oxidative stress responses, altered carbon regulation, and enhanced biofilm formation (Fig. 5D). Many of these correspond to pathoadaptive loci frequently mutated during *P. aeruginosa* evolution in chronic lung infections (*46*).

Among these pathoadaptive genes, our Tn-seq results specifically suggest a prevalent role for biofilm formation in adapting to the PYO5-diff HBE mucosal surface. Biofilm-deficient mutants such as *pslA*, which cannot synthesize the polysaccharide Psl, showed lower fitness in PYO5-diff tissues. In contrast, *wspF* mutants, which produce excessive matrix leading to continuous biofilm formation (*47*), show increased fitness in PYO5-diff compared to untreated tissue (Fig. 5E). Hyperbiofilm mutants, such as those with loss-of-function in *wspF*, often arise during chronic infections (*12*, *46*). We have previously reported a notable fitness tradeoff for these mutants when growing on an untreated mucosal surface: they show increased recalcitrance to antibiotic treatment but pay high metabolic and colonization costs in the absence of drug selection (*45*). Our results suggest that PYO-induced tissue remodeling could at least partially alleviate the bacterial growth impairments caused by these mutations, helping them regain higher fitness levels. This motivated us to directly test such a prediction in our model.

### Remodeled mucosa increases the fitness of hyperbiofilm mutants

To directly test whether PYO-remodeled tissue reduces the fitness costs associated with forming biofilms, we used an imaging approach and infected tissues with the hyperbiofilm-producing Δ*wspF* strain constitutively expressing mNeonGreen. The mutant grew more extensively in PYO-diff tissues relative to untreated controls, forming large biofilm aggregates (Fig. 6A-B). Biofilms seemed to preferentially accumulate in squamous regions adjacent to thicker tissue (Fig. S7), the same regions we found showed disrupted ciliation (Fig. 3C, G, J, K). Locally increased biofilm growth eventually led to greater tissue damage compared to untreated controls (Fig. 6B-D). Fluorescence and CFU quantification confirmed that Δ*wspF* grew more extensively in PYO5-diff tissues (Fig. 6C, E). Notably, Δ*wspF* showed a greater fitness advantage on PYO5-diff tissues compared to NC than the wild type, indicating it may benefit more under these conditions (Fig. 6F).

**Figure 6.**
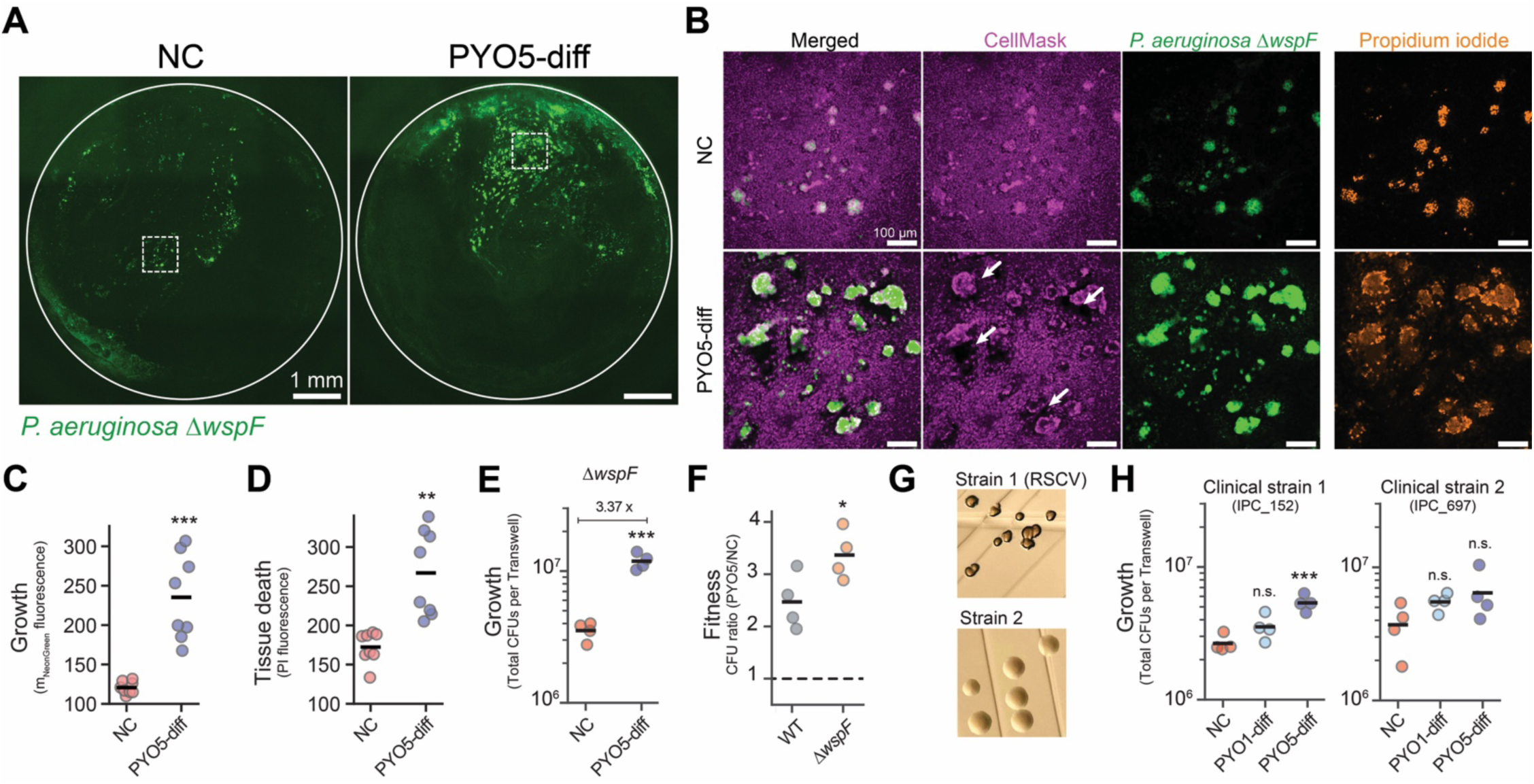
Remodeled mucosa increases the fitness of hyperbiofilm mutants. **A.** Representative images of *P. aeruginosa* Δ*wspF* growth 9-10 hours post-infection in airway epithelia differentiated with and without chronic exposure to PYO. Circles represent the entire tissue area, and dotted squares highlight zoomed-in areas shown in panel B. **B.** Representative zoomed in areas showing Δ*wspF* growth at the mucosal surface and its resulting tissue damage, visualized with propidium iodide. Arrows in the CellMask channel point to some of these heavily damaged areas in the tissue (see Movie S10-11 for full z-stacks). **C-D**. Quantification of *P. aeruginosa* Δ*wspF* growth (C) and inflicted damage (D) based on fluorescence signal during live microscopy after 9-10 hours post infection. Each data point represents one field of view (n = 8) among three biological replicates; horizontal black lines mark their mean. **E.** Quantification of *P. aeruginosa* Δ*wspF* growth with CFUs plated immediately after images were collected. Each data point represents an independent biological replicate (n = 4); horizontal black lines mark their mean. **F.** Comparison of fitness in PYO5-diff vs NC for WT and Δ*wspF* strains. Fitness is represented by the CFU ratio in PYO5-diff/NC, with values above 1 indicating increased growth in PYO5-diff tissue. Values for WT are derived from data in Fig. 1J, while values for Δ*wspF* are derived from data in Fig 6E. Each data point represents an independent biological replicate (n = 4); horizontal black lines mark their mean. **G.** Colony morphology of the two clinical strains containing *wspF* mutations used in the study. Note the dramatic RSCV phenotype of strain 1, a characteristic of strains overproducing extracellular polysaccharides. **H.** Quantification growth of the two *P. aeruginosa* clinical strains in tissues chronically exposed to distinct concentrations of PYO. Growth was assessed by CFUs platting after 9-10 hours post-infection. Statistics in **C-F**, Welch unpaired t-test (two-sided); in **H**, one-way ANOVA with Tukey HSD multiple comparison test, with asterisks showing significant differences relative to NC (\**p* < 0.05; \*\**p* < 0.01; \*\*\**p* < 0.001; n.s., *p* > 0.05).

Finally, since many of the genes selected for in PYO-diff tissues are linked to pathoadaptation (*46*), we wondered whether epithelial remodeling could improve the fitness of strains sampled from chronically infected patients. We obtained clinical isolates carrying *wspF* mutations that evolved during human chronic infection: one isolate displayed a rugose small-colony variant (RSCV) phenotype, a hallmark of hyperbiofilm strains, and the other exhibited a regular colony morphology (Fig. 6G). Despite both isolates showing a trend toward increased growth in PYO-diff tissues, the RSCV strain showed the greatest increase in fitness, consistent with PYO-diff tissues alleviating the burden of hyperbiofilm formation (Fig. 6H).

Overall, these results demonstrate that PYO exposure during differentiation generates a distinct tissue microenvironment that reshapes selective pressures, favors pathoadaptive traits, and mitigates some of the fitness costs associated with mutations commonly acquired during chronic infection.

## Discussion

Our study demonstrates that exposure to the bacterial metabolite PYO disrupts airway regeneration by inducing epithelial defects that impair mucociliary clearance. This disruption creates a permissive environment for chronic *P. aeruginosa* infection, specifically favoring the growth of mutants commonly isolated from clinical contexts, particularly those exhibiting enhanced biofilm formation. Collectively, these findings illustrate a feedback loop wherein microbial metabolites remodel host tissues and actively reinforce pathogen adaptation.

Hyperbiofilm strains are substantially more recalcitrant to antibiotic treatment and are recognized as primary drivers of chronicity in patients with obstructive lung diseases (*12*, *46*, *48*, *49*). The reduction of fitness costs in laboratory and clinical hyperbiofilm strains within a PYO-remodeled epithelium indicates that the specific mucosal environment plays a crucial role in determining mutant fitness and colonization. While longitudinal clinical data from chronically infected patients have long indicated this scenario as infection progresses (*12*, *48*, *49*), to the best of our knowledge, our study is the first to recapitulate these host-mediated effects under controlled experimental conditions, highlighting the advantages of using physiologically relevant human tissue culture models (*50*, *51*). This remodeled tissue-driven selection may act synergistically with antibiotic pressure to further accelerate the evolution of these lineages. The mechanisms behind this increased fitness for biofilm formation remain to be determined and may include differences in mucus composition, oxygen gradients, and different nutrient availability. Finally, given that PYO is only one of many metabolites produced by *P. aeruginosa* during infection, alongside other molecules such as distinct phenazines, quorum-sensing molecules, and siderophores (*4*, *5*, *7*), our findings provide a framework for exploring how a broader “metabolome” modulates interactions between bacteria and the epithelium. Such broader future characterization will allow distinction of effects specific to certain metabolites from effects induced by a class of molecules that can be predicted (e.g., redox activity and oxidative stress). Importantly, regardless of the specific metabolite, our findings suggest that *P. aeruginosa* does not merely damage the airway epithelium during infection but actively reshapes it, with likely consequences for both pathogen evolution and the tissue’s capacity to recover.

Inflammation has long been recognized as a central factor in infection progression. By contrast, the mechanisms by which infectious agents directly perturb airway recovery have received far less attention (*52*). The squamous expansion observed in our regenerating HBE cultures recapitulates key features of chronic airway diseases such as COPD (*53*), suggesting that microbes can actively reshape the environment to potentiate chronicity. Specifically, we identified a link between PYO exposure during differentiation and the expansion of squamous cells that may resemble recently described airway hillocks (*39*), whose functions in human airways remain largely unclear. Although these squamous cells may exhibit increased tolerance to PYO toxicity, their expansion, combined with abnormal ciliation in the same regions, creates cilia-deprived niches that promote biofilm formation and leave the tissue vulnerable to recurrent damage. PYO-induced perturbations could thus fuel a vicious cycle of infection (*26*, *27*) that may sustain long-term, detrimental interactions between microbes and airway tissue (Fig. 1K, Fig. 7).

**Figure 7.**
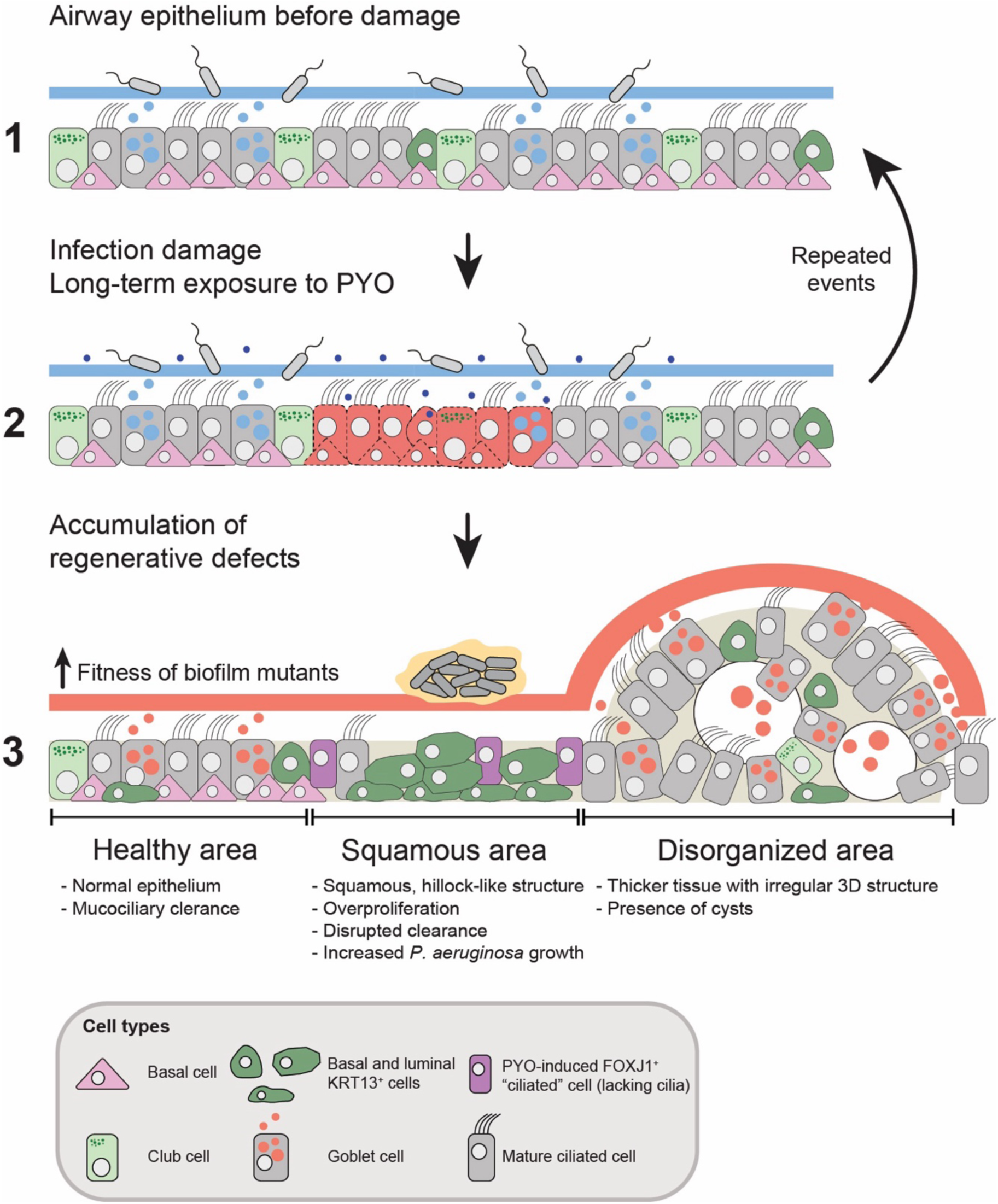
Working model of the effect of PYO in the airway epithelium. A vicious cycle of long-term exposure to PYO and infections may disrupt regeneration, leading to an altered epithelium with impaired physiological function. The altered epithelium exhibits heterogeneous spatial characteristics, including healthy (i.e., similar to untreated controls), squamous (characterized by expansion of squamous KRT13^+^ cells and cilia depletion), and disorganized areas (thicker tissue mounds with cysts). Such heterogeneity increases susceptibility to new infection damage, further supports *P. aeruginosa* growth, and facilitates the better establishment of chronically associated hyperbiofilm mutants.

Despite capturing the effects of molecules like PYO over the differentiation timescale (i.e., several weeks), our model does not capture the entire process of tissue remodeling that occurs *in vivo* during chronic *P. aeruginosa* infections. In these patients, infections can last for years, and exposure to metabolites like PYO is likely heterogeneous over time. In addition, during this exposure, epithelial cells interact with components of the immune system, and the extracellular matrix undergoes remodeling (*54*, *55*). However, the phenotypes and processes identified in human *in vitro* models, such as those we observed with PYO, provide clear targets for cellular states and gene markers that can be investigated more directly in complex human clinical samples.

More broadly, our work provides direct evidence that sustained pathogen-host crosstalk can modulate tissue development over time. To date, most research on this topic has focused on the gut, where microbial metabolites influence epithelial and immune cell differentiation, as well as epithelial regeneration (*56–60*). For example, a recent study has shown that microbiome-derived nicotinic acid interacts directly with the gut epithelium to establish colonocyte identity (*61*). Similarly, distinct bacterial metabolites and toxins are known to modulate inflammation and host cell metabolism within the intestinal landscape (*59*, *62*). In the lung, however, the long-term effects of diverse secreted molecular factors (*4–7*) on tissue physiology, particularly during regeneration, remain largely unexplored. Our findings support a model of microbial pathogenesis in which secreted metabolites serve not only as acute virulence factors but also as long-term modulators of host tissue physiology by interfering with regeneration. Targeting these processes in the future could offer alternative treatment strategies for patients with chronic infections.

## Acknowledgments

We thank the Gene Expression Core Facility at EPFL and the Lausanne Genomic Technologies Facility at UNIL for sequencing (scRNA-seq and Tn-seq, respectively). We also thank the EPFL Histology Facility and the EPFL BioImaging and Optics Platform (BIOP) for assistance with histology preparation and imaging, respectively. We thank the Persat lab for constructive feedback throughout the project’s development, Andres Floto and Aaron Weimann (The University of Cambridge) for providing the clinical strains, and Dianne K. Newman for providing the PA14 strains. Finally, we thank John Ciemniecki, Juan Hernandez Bird, Bart Deplancke, Vincent Gardeux, Jayaraj Rajagopal, Jiawei Sun, and Viral Shah for their comments on the manuscript. ChatGPT-5 was used to polish grammar and syntax in this manuscript.

## Funding supporting this work

- Swiss National Science Foundation (SNSF): 310030_189084 (AP); NCCR AntiResist (AP); 200021_188623 (VH), and NCCR Microbiomes grant numbers 180575 and 225148 (VH).
- European Molecular Biology Organization (EMBO) Postdoctoral Fellowship ALTF 12-2022 (LAM).
- Cystic Fibrosis Foundation: PERSAT24I0
- EPFL Summer Research Program funding for AG

## Contributions

Conceptualization: L.A.M. and A.P.

Data curation: L.A.M., A.G., L.L.B., D.L., and E.V.

Formal analysis: L.A.M., A.G., L.L.B., D.L., and E.V.

Funding acquisition: L.A.M, V.H., and A.P.

Investigation: L.A.M., A.P., L.L.B., D.L., and E.V.

Methodology: L.A.M., A.G., L.L.B., D.L., E.V., T.D., Z.A.;

Project administration: L.A.M. and A.P.

Supervision: V.H. and A.P.

Visualization: L.A.M., A.G., L.L.B., D.L., and E.V.

Writing-original draft: L.A.M. and A.P.

Writing-review and editing: L.A.M., A.G., L.L.B., D.L., E.V., A.P.

## Methods

### General bacterial culturing

Bacteria were cultured using LB broth and agar (1.5%), supplemented with antibiotics when necessary. The antibiotics were prepared as concentrated stock solutions (100x or greater) and stored at-20 °C until use. Unless specified, all liquid cultures were incubated at 37 °C with shaking at 225 RPM (revolutions per minute).

### Bacterial strain construction

We used distinct strains of *P. aeruginosa* in our experiments. Most of the experiments were performed with PAO1 and its derivatives, but we also used PA14 and two clinical strains, as specified in the legends. A complete list of strains, plasmids, and primers used is provided in Table S10.

### Tissue culturing for Transwell HBE cultures

HBE cells were obtained from Lonza (Cat#: CC-2540, Lot#: 22TL332331 for healthy donor; Cat#: PN-00196979, Lot#: 450918 for the CF donor). Culturing followed the protocols we recently described (*45*, *64*). In summary, cells were cultured in T-25 flasks using PneumaCult-Ex Plus medium (P-Ex Plus) (Stemcell Technologies) for expansion and stored in liquid nitrogen at-180 °C for further use, always limiting expansion of cells to a maximum of three passages. We used cells from the CF donor in the first experiments to test if PYO-induced phenotypes were reproducible in cells from different donors (Figs. 1E and S2A). All other experiments used cells from the healthy donor. Incubations were always done at 37 °C with 5% CO2 using a humidified cell culture incubator. After reaching confluence in T-25 flasks, the cells were detached using the Animal Component-Free Cell Dissociation Kit (Stemcell Technologies), centrifuged, resuspended in P-Ex Plus, and loaded into Transwells. However, we performed the adaptations described below.

During culturing in Transwells, HBE cells were loaded on tissue-culturing inserts (0.4 µm pore) of polyethylene terephthalate with transparent membranes (Sarstedt). We loaded 30,000-50,000 cells per insert, which was followed by an expansion phase with P-Ex Plus until reaching confluence (100 µL on apical side, 500 µL on basal side, 3-5 days) and a transition to ALI culture conditions. ALI culturing contained PneumaCult-ALI medium (P-ALI) (Stemcell Technologies) on the basal side (500 µL) and air on the apical side. Cultures were allowed to stabilize at the ALI for 2-3 days, after which PYO treatment or control conditions were initiated. PYO was added to the basal side at a final concentration of 1-5 µM, as specified in the experiment. 10 mM PYO stocks were made in 20 mM HCl solution that was then diluted in water to 100x stocks (100-500 µM PYO), which were added to the HBE cultures. The same dilution and volumes were used for 20 mM HCl in the control, and these volumes did not alter the pH of the HBE culturing medium. Culturing media (P-ALI + PYO or P-ALI + control) were replaced in the basal side 3x a week, and differentiation proceeded 4-6 weeks, when the cultures were used as described in each experiment. Importantly, PYO was added to the basal side for technical reasons, as HBE cells are stressed when liquid is added to the apical (air) side, preventing controlled long-term apical exposure.

### Single-cell RNA-seq (scRNA-seq)

#### Experimental design

HBE cultures were prepared as described above and kept under differentiation for 36 days. Our scRNA-seq experiment consisted of four conditions: 1. Negative control (NC): No PYO added, HCl added as a negative control following the exact amount added in the PYO treatments; 2. PYO shock: 1 µM of PYO was added ∼ 24h before cells were collected; 3. PYO1-diff: 1 µM PYO kept throughout the differentiation; PYO5-diff: 5 µM of PYO kept throughout the differentiation (Fig. 2A).

#### Sample processing

Sample processing followed the protocols previously described (*64*), with modifications. In summary, HBE cultures were first washed twice with PBS, the membranes were cut with a sterile scalpel, and placed into 1.5 mL of dissociation buffer with the following composition: 300 µL Protease from *Bacillus licheniformis* (100 mg/ml, Sigma), 3 μL DNase I (10 mg/mL, Roche), 30 μL EDTA (0.5 M, Sigma), 30 μL EGTA (0.5 M, BioWorld), 237 μL sterile PBS, and 900 μL Accumax (Brunschwig). Four HBE cultures were used for each treatment, and dissociation of each culture was performed separately. Epithelia were incubated in dissociation buffer at 37 °C for 5 min and then mixed with a 1000 µL and 200 µL pipette (25-50 times with each). This process was repeated if cells were still not well dissociated. Then, membranes were removed from each sample, followed by centrifugation for 5 min at room temperature (RT) at 400xg. Supernatant was removed, leaving a residual volume of ∼ 50 µL. This pellet was mixed in the residual volume using a 200 µL pipette. Then, cell suspensions from the four HBE cultures for each treatment were combined and topped up with 1 mL of pre-chilled 0.04% molecular-grade BSA in PBS. This entire dissociation process took around 30 minutes. From this point, all steps were performed on ice or at 4°C. Residual mucus clumps floating in the sample were removed. Then, each sample was filtered through a 40 μm Flowmi cell strainer (Bel-Art) and centrifuged (10 min, 4 °C, 400xg). Next, fixation was processed according to 10X Genomics recommendations. For each sample, the supernatant was carefully removed, and cells were resuspended in 1 mL of 10X Genomics fixation buffer (791.9 µL of nuclease-free water, 100 µL of Conc. Fix & Perm Buffer, 108.1 µL of molecular grade formaldehyde 37%). Fixation was performed for 22 h at 4 °C without agitation. Samples were then centrifuged (5 min, RT, 850xg), supernatants were removed, and pellets were re-suspended in 1 mL of pre-chilled Quenching Buffer (875 µL of nuclease-free water, 125 µL of 8x Conc Quenching buffer). Finally, 0.1 volume of pre-warmed Enhancer (10x Genomics PN-2000482) was added to each fixed sample in Quenching buffer, followed by the addition 50% glycerol to a final concentration of 10%. Aliquots were used for cell counting, and samples were stored at-80 °C until proceeding for sequencing.

#### 10x Genomics Flex single-cell gene expression sequencing

Sequencing was done at the EPFL Gene Expression Core Facility (GECF). In summary, fixed cells were thawed and washed with the Chromium Next GEM Single Cell Fixed RNA Sample Preparation Kit (PN-1000414, 10x Genomics, Pleasanton, CA), according to the manufacturer protocol CG000478, Rev D. Probes hybridization with barcodes for multiplexing (Chromium Human Transcriptome Probe Set v1.0.1), as well as cells pooling and washing were performed before loading into a Chromium X Single Cell Controller (10x Genomics) in a chip together with beads, master mix reagents and oil to generate single-cell-containing droplets. Four samples were pooled, and 4,000 cells per sample were targeted for a total of 16,000 cells in the pool. Cell counts and quality were assessed at several stages using the Trypan blue method before pooling and loading. Finally, single-cell Gene Expression libraries were prepared using the Chromium Fixed RNA Profiling Kit (10x Genomics), according to the manufacturer’s protocol CG000527 Rev E.

The library’s quality control was performed using a TapeStation 4200 (Agilent) and a Qubit dsDNA High Sensitivity Assay (Thermo) according to the manufacturer’s instructions. The sequencing libraries were loaded on an Aviti Cloudbreak Freestyle (Element Biosciences) flow cell and sequenced in HD mode with the manufacturer’s short inserts recipe, with a read1 of 29nt and a read2 of 91nt, at a mean depth of about 35k reads/cell. Bases2fastq v1.7.0 (Element Biosciences) was used to demultiplex reads, which were then quality-controlled with fastQC v0.11.9. FastQ Screen v0.14.0 was used for screening FASTQ file reads against multiple reference genomes.

Cell Ranger Single Cell Software Suite v8.0.0 from 10x Genomics was then used to perform sample demultiplexing and probe counting using 10X Genomics custom annotation of human genome assembly GRCh38-2024-A.

#### Data pre-processing

Low-quality cells were filtered out based on the percentage of mitochondrial reads, the number of genes, and transcripts detected per cell. Cells with higher than 25% of mitochondrial reads and with fewer than 1000 detected genes were considered dead or empty droplets. Also, outlier cells from each sample were deleted. Each sample was processed separately using the Seurat R package and the SCT data transformation, with default parameters. Doublet Cells were identified and removed using the DoubletFinder R package (*67*). Subsequently, all samples were integrated using the updated Seurat integration pipeline for SCT-transformed data. PCA was performed for linear dimensionality reduction, and the top 20 PCs were selected for further analysis. This selection was guided by visual inspection of PC heatmaps and by assessing the biological relevance of genes associated with the principal components. The unsupervised Louvain algorithm initially clustered cells into 11 clusters at a resolution of 0.5.

#### Clusters characterization

For cluster characterization and cell type identification, marker genes for canonical airway epithelium cell types were collected from the literature (*69*, *70*) and the Human Protein Atlas (https://www.proteinatlas.org/) (*71*). Genes that had variable expression in our dataset were selected and used further for semi-supervised cell type annotation using scSorter (*72*). In the first round of annotation, we identified broader cell types: basal, secretory, ciliated, and ionocyte. A second step consisted of the identification of more granular, specialized cell types, such as hillock-like, club, and goblet cells, based on expression of KRT13, SCGB1A1, and MUC5B, respectively. Based on the expression of canonical markers, some major clusters were separated and sub-clustered with a resolution of 0.2. Clusters expressing a mix of genes were annotated as transitory. Ciliated cells were separated and sub-clustered to verify the differences in PYO exposure in a specific subset. Finally, marker genes for each cluster were identified and used to manually verify the annotation. Gene markers were identified using the Wilcoxon Rank Sum test implemented in the FindAllMarkers function of the Seurat R package. The log_2_FC threshold was set to 1, the minimum percentage of cells expressing the markers was set to 0.7, and the minimum percentage of difference between the two groups was set to 0.3 to ensure identification of robust markers. Genes with adjusted p-value smaller than 0.05 were considered significant. A combinatorial score was given to a gene based on rankings of the following parameters: log2FC, percentage of cells expressing the gene in the cluster of interest, and percentage of cells expressing the gene outside the cluster of interest. Genes with the highest scores per cluster were considered to be the most descriptive ones. Those were annotated using the biomaRt R package (*73*) and used for manual cell-type verification.

#### Differential gene expression (DGE) analysis between ciliated clusters

First, differentially expressed genes in ciliated clusters 15 versus 14 of the PYO5-diff sample were computed using the Wilcoxon Rank Sum test implemented in the FindMarkers function of the Seurat R package with default parameters. Genes with adjusted p-value smaller than 0.05 were considered significant and selected for differential gene set enrichment. Differentially expressed genes later were divided into up-and down-expressed genes (in cluster 15 vs cluster 14) and used for Gene Ontology Enrichment via the enrichGO function of the clusterProfiler R package (*74*), with default parameters. Terms with adjusted p-value lower than 0.05 were considered significant.

#### DGE for metabolic and signaling analyses

For each cluster, differentially expressed genes across 3 conditions (Shock, PYO1-diff, PYO5-diff) relative to the control were identified using the FindMarkers function in the Seurat R package. The minimum percentage difference between two groups was set to 0.3, and the remaining parameters were kept at their defaults. Genes with adjusted p-value smaller than 0.05 were selected for signaling and metabolic analyses. Differentially expressed genes were also identified using a pseudo-bulk approach to increase statistical power and inform the metabolic modeling (see next). In this case, conditions were compared to the control regardless of clusters, along with the default parameters of the FindMarkers function. Genes with adjusted p-value smaller than 0.05 were selected for signaling and metabolic analyses.

### Metabolic modeling

#### Reconstruction of a metabolic model for HBE cells

To investigate the metabolic effects of PYO in HBECs, we reconstructed a reduced, context-specific metabolic model focused on subsystems of interest. Model reduction was performed using the redHUMAN algorithm (*75*), starting from the human genome-scale metabolic network Recon3D (*76*). The extracellular environment was defined according to the composition of DMEM.

A total of 30 metabolic subsystems were selected as starting points for model reconstruction, including glycolysis/gluconeogenesis, the citric acid cycle, pentose phosphate pathway, reactive oxygen species (ROS) detoxification, arachidonic acid metabolism, eicosanoid metabolism, keratan sulfate degradation and synthesis, sphingolipid and glycosphingolipid metabolism, linoleate metabolism, lysine metabolism, O-glycan metabolism, leukotriene metabolism, N-glycan degradation, coenzyme A catabolism, vitamin C metabolism, androgen and estrogen synthesis and metabolism, chondroitin sulfate degradation, cholesterol metabolism, NAD metabolism, heme degradation, tryptophan metabolism, vitamin A metabolism, fatty acid synthesis, pyruvate metabolism, folate metabolism, glutathione metabolism, inositol phosphate metabolism, and oxidative phosphorylation.

Subsystem selection was guided by enrichment analysis, focusing on (i) the most enriched metabolic subsystems in the negative control condition, capturing the characteristic metabolic signature of HBECs, and (ii) the most deregulated subsystems across the three PYO treatment conditions, capturing metabolic alterations induced by PYO exposure.

For each selected subsystem, a corresponding set of metabolic tasks was defined. Model reduction was performed using redHUMAN with default parameters, setting D = 1 for redGEM and using the Smin option for lumpGEM. To ensure that the reduced model retained the ability to carry out all specified metabolic functions, lumpGEM was applied independently for each metabolic task, in addition to biomass production.

The resulting model, termed redHBECPYO, comprises 1,004 unique metabolites across cellular compartments, 1,458 genes, and 4,127 reactions, spanning 71 metabolic subsystems.

#### Integration of transcriptomic data and estimation of representative fluxes

We first identified metabolic genes present in the redHBECPYO model that were detected in the pseudo-bulk RNA-seq dataset. Differential expression analysis was used to classify genes as upregulated or downregulated using a fold-change threshold of 1.5 across all pairwise comparisons between conditions (i.e., each treatment compared to the negative control or to another treatment). Genes with a fold change greater than 1.5 were considered upregulated, whereas genes with a fold change lower than 0.66 were considered downregulated in at least one comparison.

Using the gene-protein-reaction (GPR) rules encoded in the metabolic model, gene expression changes were mapped to the corresponding metabolic reactions. In total, 912 genes passed the fold-change threshold, encoding 1,404 reactions and yielding 3,677 flux ratios, each defined between a treatment condition and either the negative control or another treatment.

To integrate the transcriptomic data into the redHBECPYO model, we applied the REMI method (*77*), which assumes that relative changes in gene expression translate into proportional changes in enzyme abundance and, consequently, reaction rates. REMI constrains the metabolic network such that reactions associated with upregulated genes are forced to carry higher fluxes, whereas reactions associated with downregulated genes are constrained to carry lower fluxes. In this study, we used an extended version of REMI that allows the simultaneous integration of more than two conditions by incorporating the corresponding pairwise flux ratios across all conditions. The method then identifies a flux distribution that maximizes the number of reaction rate constraints that can be satisfied simultaneously for each pairwise condition comparison.

REMI analysis showed that, out of the 3,677 flux ratios, a maximum of 3,589 could be simultaneously integrated into the model, indicating that the transcriptomic expression profile is largely consistent with the network mass-balance constraints. Notably, three alternative optimal solutions were identified, each integrating 3,589 flux ratios, of which 2,954 were common across all alternatives.

The 2,954 flux ratios common to all REMI solutions were fixed in the redHBECPYO model, and the resulting constrained solution space was sampled using the Artificial Centering Hit-and-Run (ACHR) algorithm. A total of 250,000 flux samples were generated.

For each reaction, we calculated the mean flux across samples as well as the mean fold change between each PYO treatment and the negative control. These mean values were used as representative reaction fluxes for all downstream analyses.

#### Minimal network enrichment analysis to study metabolic functions

To investigate the effects of PYO on metabolic functions in HBEC cells, we defined metabolic tasks informed by pseudo-bulk RNA-seq data, focusing on metabolic subsystems that were enriched in the negative control condition and/or deregulated across the three PYO treatment conditions (Table S1-S2).

For each metabolic task, we constructed minimal functional networks by formulating a mixed-integer linear programming (MILP) problem that identifies the smallest set of reactions required to accomplish the task, while also allowing for alternative feasible pathways (*78*). This approach yields task-specific minimal networks that capture the essential metabolic routes supporting each function.

To quantify task activity, we used the representative reaction fluxes obtained from flux sampling of the constrained redHBECPYO models. For each metabolic task, only reactions that were present in all alternative minimal networks associated with that task were considered, ensuring that the resulting activity measures reflect core, task-essential metabolic reactions.

#### Enrichment analysis of signaling pathways

To relate the metabolic changes observed under PYO treatment to upstream regulatory programs, we analyzed signaling pathways in a mechanistic network centered on transcription factors (TFs) that regulate metabolic genes. We used a TF-specific signaling network reconstructed by Liaskos, Masid et al. (unpublished), which comprises 9,227 reactions and 12,288 species organized into 268 Reactome-annotated signaling pathways (*79*) and includes 115 TFs with regulatory links to metabolic genes in Recon3D (*76*).

For each of the three PYO treatments versus the negative control, we performed pathway over-representation analysis using an upper-tailed hypergeometric test separately for upregulated and downregulated genes, using both (i) population-level and (ii) cell-type-resolved differential expression data. Based on the differential expression data, we defined upregulated genes as those with a fold change > 1.5 and an adjusted p-value < 0.05, and downregulated genes as those with a fold change < 0.66 and an adjusted p-value < 0.05. We corrected hypergeometric-test p-values using the Benjamini-Hochberg procedure across the 268 pathways for each gene set and comparison. To assign a direction of deregulation to each pathway, we compared the fraction of upregulated and downregulated genes among pathway members. We assigned a positive sign to pathways with a larger fraction of upregulated genes and a negative sign to pathways with a larger fraction of downregulated genes.

To facilitate biological interpretation within the context of human primary bronchial epithelial cells, six Reactome pathway labels were manually modified (Fig. 2D and Fig. S5D) to reflect the underlying molecular function rather than tissue-specific discovery contexts. Specifically, “Neutrophil degranulation” was labeled “Exocytosis of granule proteins”; “Bile acid and bile salt metabolism” was labeled “Extrahepatic cholesterol catabolism”; and “Cell surface interactions at the vascular wall” was labeled “Cell adhesion molecule interactions”. Additionally, three lengthy database identifiers were shortened: “Gene and protein expression by JAK-STAT signaling after Interleukin-12 stimulation” was labeled “Interleukin-12 stimulated gene expression through JAK-STAT”; “Regulation of Insulin-like Growth Factor (IGF) transport and uptake by Insulin-like Growth Factor Binding Proteins (IGFBPs)” was labeled “Regulation of IGF transport and uptake by IGFBPs”; and “HSP90 chaperone cycle for steroid hormone receptors (SHR) in the presence of ligand” was labeled “HSP90 chaperone cycle for SHR in the presence of ligand”. The original Reactome identifiers are retained in Tables S4 to ensure traceability.

#### Immunofluorescence

All HBE cultures used in immunofluorescence (IF) were kept in differentiation (with or without PYO) for 4-6 weeks, except the regeneration assays, which are described in the next session. All steps were performed at RT temperature, and all the procedures were performed in Transwell inserts, without cutting the membrane, with treatments added to the top part of the membrane. HBE cultures were first washed with PBS (5 min incubation) for the removal of mucus. Samples were then fixed with 4% paraformaldehyde (PFA, Electron Microscopy Sciences) for 15 min, washed with PBS, permeabilized with 0.2% Triton X-100 solution (VWR Life Science) for 20 min, and exposed to a blocking solution (5% BSA, 0.2% Triton in PBS) for 20 min. Subsequently, samples were incubated with primary antibody solutions for 1 h. These included rabbit anti-KRT13 (ab92551 – Abcam, 1:250), rabbit anti-alpha tubulin (ab52866 – Abcam, 1:250), rabbit anti-Ki67 (MA5-14520 – ThermoFisher, 1:250), mouse anti-FOXJ1 (14-9965-82 – eBioscience, 1:100), and mouse anti-acetylated tubulin (T7451 – Sigma-Aldrich, 1:250). Samples were then washed with PBS and incubated with secondary antibodies for 1 h in the dark. Secondary antibodies included goat anti-rabbit IgG H&L Alexa Fluor 488 (ab150077 – Abcam, 1:200) and goat anti-mouse IgG H&L Alexa Fluor 594 (A-11005 – ThermoFisher, 1:200). Finally, nuclei and actin were stained with DAPI (Sigma-Aldrich, 1:1000) and Phalloidin Atto 655 (18846 - ThermoFisher, 1:100), respectively, for 10 min.

Imaging of fixed samples was performed with a Nikon Eclipse Ti2-E inverted microscope coupled with a Yokogawa CSU W2 confocal spinning disk unit. We used a mix of 4x, 20x, and 40x objectives (Nikon MRL00042, MRD77200, and MRD77410, respectively). Imaging at 4x was initially used to characterize the overall tissue structure, followed by z-stacks of zoomed-in areas at 20x or 40x. Details on imaging for each sample are described in the “imaging and image analysis session”.

#### Regeneration assays

For regeneration assays (displayed in Figs. 1A-B and Fig. S1), HBE cultures were first differentiated without any treatment for 5 weeks, when mucus was washed from the cultures with PBS, followed by one day of stabilization. HBE cultures were then damaged with a pipette tip in an “X” format and then incubated with or without PYO (5 µM) during their recovery for the next 30 days. Additional cultures from the same batch were used as “no damage” controls. The protocol was the same, except for the damage. Three replicates were prepared for each condition. After 30 days of recovery, samples were imaged in bright-field mode, fixed, and processed for immunofluorescence as described above.

#### Histological analyses

HBE cultures used for histology were differentiated with and without PYO (5 µM) for 30 days. We prepared two samples for each treatment. HBE cultures were washed three times with PBS to remove mucus and fixed from the top and bottom of the insert with 4% PFA in PBS overnight at 4 °C. Then, samples were washed again with PBS, and the Transwell insert membranes were cut with a scalpel, stored in PBS, and processed for mounting, sectioning, and H&E staining by the EPFL Histology Core Facility using standard histology protocols. Slides were then imaged with a Leica SP8 upright microscope at the BioImaging and Optics Platform (BIOP).

#### Mucociliary clearance assay

HBE cultures used for mucociliary clearance were differentiated with and without PYO (5 µM) for 35 days. Three biological replicates (i.e., Transwell inserts) were used for each treatment. The apical part of these cultures was first washed with PBS, and then filled with 50 µL of a 1:250 solution of yellow-green carboxylated fluorescent beads with a 2 μm diameter (FluoSpheres, Life Technologies). Inserts were then transferred to custom-designed PDMS inserts, which were plasma-bonded to the glass bottoms of 12-well plates. P-ALI medium was used to fill the bottom of the culture, and the plate was incubated in a UNO-T-H-CO2 stage-top incubator (Okolab) under the same conditions (37 °C with 5% CO2 and humidity). Cultures were incubated for ∼30 min under these conditions to stabilize, and imaging began immediately thereafter. Bead flow was imaged with our spinning disk confocal microscope described above in the GFP channel. Focus was adjusted to ∼75 µm above the tissue with a 4x objective, and images were acquired at 0.25 s intervals.

### Imaging and image analyses

#### Particle Image Velocimetry (PIV)

Velocity vector fields and velocity distributions were quantified from time-lapse image sequences of 2 µm-wide fluorescent bead motion acquired ∼75 µm above the tissue using a 4x objective (Nikon MRL00042) at 0.25 s intervals, using a custom Python pipeline (Fig. 1H and Fig. S3). Each pair of successive frames was first preprocessed by subtracting their mean intensity to suppress static features (e.g., beads adhered to mucus strands). Image stacks were then divided into interrogation windows of 32 x 32 pixels with 50 % overlap, and velocity vectors were computed using an FFT-based cross-correlation approach with a 1.2x enlarged search area using the *openpiv* package. Spurious vectors were removed using a signal-to-noise ratio threshold of 1.1 and replaced for up to 3 iterations with local mean values using a 3×3 kernel. For each movie, the instantaneous velocity components (u, v) and their spatial coordinates (x, y) were computed for all frame pairs. Velocity field maps were generated by taking the median u and v components across the full sequence. Velocity colormaps were generated by computing the temporal median of the velocity magnitude and displayed using the viridis colormap. The distribution of velocity values was exported as CSV files for quantitative comparison across datasets. This data was used to plot the distributions shown in Fig. 1I.

#### Surface coverage

KRT13^+^ and ciliated (AcTub^+^) cells imaged with a 4x objective (Nikon MRL00042) were segmented by applying a fixed intensity threshold (250 and 190, respectively) to generate binary masks, where pixels above the threshold were considered positive for staining. For both stainings, the coverage of the Transwell surface was then quantified by calculating the fraction of positive pixels included in the total Transwell area and expressed as percentage. The Transwell area was computed based on the known Transwell diameter (6.5 mm). To quantify the spatial relationship between the two cell types, the binary masks of the AcTub and KRT13 stainings were combined using a logical XOR operation, thereby identifying pixels that were positive for one marker but not both. The anticorrelation index was then defined as the number of XOR-positive pixels divided by the sum of the AcTub and KRT13 positive pixels. Plots related to these analyses are shown in Fig. 3H-I.

#### Heatmaps

Heatmaps were generated to quantify the spatial distribution of KRT13^+^ and proliferative (Ki67^+^) cells. Because Ki67 staining appears as discrete spots, spot detection was first performed in TrackMate7 using the LoG detector (object diameter: 10 µm, quality threshold: 12), and the resulting center coordinates were then used to create a binary mask. For both markers, binary masks were divided into 50×50 pixel tiles, and the local density of positive pixels per tile was computed as the fraction of positive pixels normalized by tile area (in mm^2^). Heatmaps were visualized using the *inferno* colormap and saved as TIFF images. Density values for tiles within the circular Transwell area were exported to CSV files and used to generate kernel density estimates for each replicate, with patchy spatial distributions indicated by simultaneous enrichment in low and high densities. Plots related to these analyses are shown in Fig. 3E-F and Fig. 4A-B.

#### Quantification of FOXJ1^+^ cilia^-^ cells

HBE cultures used for FOXJ1 and α-tubulin staining were differentiated with and without PYO (5 µM) for 32 days before fixation and the IF protocol described above. IF images were acquired using our 40x water-immersion objective (Nikon MRD77410). Three biological replicates (i.e., Transwell inserts) were imaged for each treatment, with four distinct areas imaged within each replicate. In each imaged area, a z-stack from bottom to apical side was acquired. Then, z-stacks were processed with Fiji. For each z-stack, a maximum intensity projection was created, and the number of cells containing FOXJ1 signal that did not display cilia staining within or next to it was manually counted. α-tubulin staining was used as a proxy for cilia here due to the availability and compatibility of the FOXJ1 antibody for dual imaging. Despite a stronger background than the cilia-specific marker we used in other experiments (acetylated-α-tubulin), α-tubulin staining was sufficient to clearly visualize cilia structures. Plots related to these analyses are shown in Figs. 3C-D.

#### KRT13 signal throughout z-depth

HBE cultures used for KRT13 and acetylated-α-tubulin were differentiated with and without PYO (5 µM) for 30-32 days before fixation and the IF protocol described above. IF images were acquired using our 20x water-immersion objective (Nikon MRD77200). Three biological replicates (i.e., Transwell inserts) were imaged for each treatment, with three to five distinct areas imaged within each replicate. All images were used for the NC, while only images of altered ciliated areas were used in the PYO5. In each imaged area, a z-stack from bottom to apical side was acquired (25 µm range for NC, 20 µm range for PYO5). Then, z-stacks were processed with Fiji. For each z-stack, the mean fluorescence intensity was measured in each z-position. These measurements and their representative images are shown in Figs. 3K-L

#### *P. aeruginosa* Δ*wspF* growth and tissue damage

HBE cultures differentiated with and without PYO (5 µM) were infected with *P. aeruginosa* Δ*wspF* expressing mNeonGreen following the protocols described below (see “Infection experiments with distinct strains” session). Propidium iodide (PI) was added to detect cell death in the tissue. After 9h of infection, Transwell inserts were imaged live using our 4x and 20x (water-immersion) objectives (Nikon MRL00042 and MRD77200, respectively). For quantification, a total of four biological replicates (i.e., Transwell inserts) were imaged. Imaging at 4x enabled characterization of the entire tissue, while 20x was used for zoomed-in areas and quantification. Imaging at 20x included acquisition of z-stacks of the entire tissue and mucosal surface, with two fields of view per biological replicate. We imaged only in areas where growth had been detected with the 4x (see Fig. 6A-B for one example). Then, z-stacks were processed with Fiji. For each z-stack, an average intensity projection was created for both the mNeonGreen and the PI channels. The mean fluorescence intensity for each image was then quantified. These measurements are shown in Fig. 6C-D.

### Tn-seq design and sequencing

#### Experimental design

Our Tn-seq experimental design followed the design we recently described (*45*), with some modifications. For the reader’s convenience, all the details are described below. We used the same Tn-library we had used before (*45*). The library stock was diluted in LB to an OD600 of 0.25 (3 ml) and grown for 14 h until deep stationary phase. Starting this culture at a high OD600 was important to limit the number of generations and consequent pre-selection before the experiment started (∼4 generations for the bulk library), while maintaining a high transposon saturation in the inoculum (*45*). Then, cells were spun down, washed in PBS, resuspended in PBS to an OD600 of 1, and used as an inoculum for HBE infections. Mucosal epithelia developed from HBE cells differentiated with and without PYO (0, 1, or 5 µM; NC, PYO1-diff, and PYO5-diff, respectively) for 45 days were used in these experiments. PYO was washed away before infection with the transposon library. Ten different Transwell inserts for each donor were infected with 1 µl, corresponding to ∼0.5-1 x 10^6^ *P. aeruginosa* cells per insert. The infection progressed for 9.5 h, resulting in ∼4 generations, with minor tissue damage in the PYO5-diff treatment and none in the other treatments. After infection, the tissue was homogenized using 100 µl of Triton X-100 (0.1% in PBS). Each insert was vigorously scraped until all the tissue material was removed. Then, samples from the ten distinct inserts were pooled, vortexed, vigorously pipetted, spun down (2 min, 14,000 RPM), washed in PBS to remove traces of Triton, spun down again, and the pellets were stored at-80 °C until subsequent processing. To determine the exact inoculum size and number of generations during infection, the inoculum and an aliquot of the stored pellet were plated for CFU.

In parallel, we prepared an LB control sample and a P-ALI control sample (Fig. 5A). The inoculum used for the infections was grown in LB or P-ALI for the same duration as the infections. In these, the inoculum was diluted to an OD600 of 0.25 in 3 ml. These cultures were grown in parallel with the infections for 9.5 h. Then, 1 ml of each culture was spun down, resuspended in Triton X-100 (0.1% in PBS), and incubated for 15 min, spun down, washed in PBS, and spun down again, and the pellet was stored at-80 °C until subsequent processing. To determine the exact inoculum size and the number of generations during growth, we plated the culture to determine CFU at the start and the end of the experiment.

This exact protocol was repeated twice on two different days to collect samples for the two biological replicates. In total, our dataset comprised 14 samples: library, inoculum, three airway mucosa (NC, PYO1-diff, PYO5-diff), and two rich media controls (P-ALI and LB), with two replicates per condition.

#### Tn-seq library preparation and sequencing

Genomic DNA (gDNA) extraction was done with the QIAamp DNA Mini kit (Qiagen). Subsequently, library preparation and sequencing were performed at the Lausanne Genomic Technologies Facility (University of Lausanne), following the protocol described previously (*45*), with minor modifications. For the reader’s convenience, the same full protocol is described below.

Overall, gDNA (500 ng) was first sheared with a Covaris S220 using 400 bp insert settings (50 µL in microTUBES with AFA fiber, Peak incident power: 175, Duty factor: 5%, Cycles per burst: 200, time: 55sec). Libraries were prepared with the xGen DNA MC Library Prep Kit (IDT, protocol version v2) using xGen UDI-UMI adapters (IDT, 15 µM stock). With these adapters, the P5 and P7 sequences are inverted compared to Illumina adapters, allowing transposon sequencing directly from read 1 (P5 side) in a single-end run (see below).

The purified ligated product was PCR amplified with a primer specific for the Illumina P7 sequence (CAAGCAGAAGACGGCATACGA) and a second one specific for the transposon sequence (cgacgttgtaaaacgaccacgt) carrying a 5’-biotin. PCR was performed with the KAPA HiFi HotStart ReadyMix kit (Roche). Cycling conditions were 95°C for 3min, followed by 12 cycles of 98°C for 15s, 60°C for 30s, and 72°C for 30s, and a final extension of 1 min at 72°C. The library was purified with SPRI beads at a 1X ratio.

The PCR product was captured with pre-washed Dynabeads MyOne Streptavidin T1 (ThermoFischer). Binding & Wash (B&W) Buffer 2X composition is 10mM Tris-HCl pH 7.5, 1mM EDTA, 2M NaCl. At least 1ml of 1X B&W Buffer was added and mixed with 25 µl of Dynabeads. After 1min on a magnet, the supernatant was removed and Dynabeads washed with the same volume of 1X B&W Buffer. This wash was repeated a second time. After removal of the supernatant on the magnet, the Dynabeads were resuspended with 50 µl of 2X B&W Buffer.

50 µl of library was mixed with the washed Dynabeads, followed by incubation at RT on a rotator for 30 min. After 2 min on a magnet and discarding the supernatant, a wash was done with 100 µl of 1X B&W Buffer. Two additional washes were done before the final elution in 40 µl H2O.

Half of the washed capture was used for the nested PCR with the Illumina P7 sequence (see sequence above) and a tailed primer made of the Illumina P5 sequence (red), the TruSeq read 1 primer binding site (black), and a transposon-specific binding sequence (orange) (for the color-coded primer, see Table S10). This nested PCR amplification was performed with the KAPA HiFi HotStart ReadyMix kit with same cycling conditions as above but 10 cycles. The final library was purified with SPRI beads at a 0.7X ratio. It was quantified using a fluorimetric method (QubIT, Thermo Scientific) and its size pattern analyzed using a fragment analyzer (Agilent).

Sequencing was performed on an Aviti (Element Biosciences) on a Cloudbreak Freestyle high-output flow cell for a 150-cycle single-end sequencing run. Clustering was performed with a 1nM library spiked with PhiX (Element Biosciences). Base calling and demultiplexing were done with bases2fastq (version 2.0).

#### Initial data processing

Sequencing read demultiplexing was performed using Bases2Fastq (v2.0; Element Biosciences). For each sample, the demultiplexed raw reads from two sequencing runs were first merged into a single file. The resulting reads were trimmed for the Tn tag using cutadapt (v.4.8, parameters:-g CCAGGACGCTACTTGTGTATAAGAGTCAG-O 20-e 0.1). Tn trimmed-reads were aligned against *Pseudomonas aeruginosa* PAO1 genome (NCBI: NC_002516) using BWA (v.0.7.18, parameters: mem-T 0-a - M). The number of reads per insertion site was computed using a custom script. Wiggle and bed files were generated for each sample (14 in total). In parallel, the number of read counts per gene locus was summarized with featureCounts (v.1.6.3, parameters:-s 0-M --fraction).

### Tn-seq analyses

#### Conditional essentiality

We used TRANSIT v3.3.13 for all analyses (*81*). “Wig” files generated in the initial data processing for each replicate were used for follow-up analyses. As described previously (*45*), due to high skewness in the data, we normalized all samples using the Beta-Geometric Correction (BGC), as recommended in the software manual (command “normalize” with-n betageom). Then, we used the Tn5 “resampling” method within TRANSIT (GUI mode) to assess the conditional essentiality of genes between two conditions in two distinct types of comparisons. First, to look for genes important in the airway mucosa, all the final samples (i.e., NC, PYO1-diff, PYO5-diff, P-ALI, and LB) were compared to the inoculum (Table S5). Then, we focused on comparisons within the airway mucosa context by comparing PYO1-diff and PYO5-diff with NC (Table S6). Within each comparison, the parameters followed default conditions (samples = 10,000, pseudocounts = 0.00, adaptive = True, histogram = False, includeZeros = True), except no normalization was used as wig files had already been normalized. For the comparison against the inoculum, we used the “adjusted p-value” < 0.05 as the cut-off for statistical significance within each comparison. Meanwhile, given that the samples were much more similar within the airway mucosa context, we used the “adjusted p-value” < 0.1 as the cut-off for statistical significance when comparing PYO1-diff and PYO5-diff to NC. In all cases, these approaches apply the Benjamini-Hochberg method for controlling the false discovery rate. Given the conservative estimates from BGC and the limited number of generations available for library growth (due to inoculum/material size), we did not restrict the results to any particular log_2_ fold-change threshold. Instead, we show all the genes that made the cut-offs in our supplementary tables (Tables S5-6). Finally, hits from PYO5-diff vs. NC conditions (adjusted p-value < 0.1) were used for enrichment analysis using the Database for Annotation, Visualization and Integrated Discovery (*82*, *83*) (https://davidbioinformatics.nih.gov), following what we have done before (*45*). We did not display the term “Two-component system” due to its lack of specificity, but the full results are available in Table S7.

#### One-way ANOVA

In addition to assessing conditional essentiality using the “resampling” method, we analyzed the variability of insertion counts across conditions in our samples using one-way analysis of variance (ANOVA) in TRANSIT. To do this, we first combined all wig files for the final samples (i.e., each replicate of NC, PYO1-diff, PYO5-diff, P-ALI, and LB) into a single wig file and applied TTR normalization. Then, this combined wig file and a metadata file were used as input for the analysis with TRANSIT, following the software’s instructions (additional parameters: normalization = nonorm, trimming = 0.0/0.0% (N/C), pseudocounts = 5, alpha = 1000.0). The results of the analysis are available in Table S8. We then selected the top 100 genes with the most significant adjusted p-values and used the seaborn.clustermap function (metric= “euclidean”, method= “average”) for visualization. The order and layout of samples within the clusters were manually adjusted to facilitate visual comparisons (Fig. 5B)

#### Fitness of disruptions of pathoadaptive genes

A recent genetics study of *P. aeruginosa* adaptation to the human lung during infections identified 224 pathoadaptive genes (i.e., genes with a mutational burden greater than expected by chance) (*46*). Because mutations in these genes are more likely to be nonsynonymous and deleterious, often resulting in protein dysfunction (*46*), we focused on how transposon-mediated disruption would affect their fitness in the airway mucosa. We first determined how many of these 224 were present in our library (i.e., non-essential genes). This reduced the number to 187. We then compared these with the genes that met our significance cutoff for the PYO5-diff vs. NC comparison (see above). This resulted in the 13 genes highlighted in Fig. 5E, which shows the fitness effects of their disruption.

### Infection experiments with distinct strains

We performed three infection experiments: (i) infections with PAO1 WT and Δ*wspF* expressing mNeonGreen (Fig. 1J; Fig. 6A-E); (ii) infections with PA14 expressing sfGFP (Fig. 1J); (iii) infections with two clinical strains (Fig. 6H) (*45*, *46*, *84*). In each experiment, strains were grown overnight in LB broth from an LB plate. Gentamicin (30 µg/mL) was added to overnight cultures of fluorescent strains. Cells were then diluted 1:100 (WT PAO1 or PA14) or 1:50 (Δ*wspF* and clinical strains) in fresh LB (no gentamicin) and incubated for 3-4 h. Cells were spun down, washed in PBS, and resuspended in PBS to an OD_600_ of 0.3, which was used as the inoculum for infections. The inoculum was plated for CFU before infection. Four Transwell inserts were infected with 1 µL of the inoculum, and infections progressed for 9-10 h. For fluorescent strains, imaging was performed after 9 h of infection, followed by CFU plating. For CFUs, the tissue was first homogenized with 0.1% Triton X-100 in PBS. Each insert was vigorously scraped until all tissue material was removed. Samples were vortexed, spun down (2 min, 12,500 RPM), and vigorously pipetted again before serial dilutions were performed in 96-well plates. We found that centrifugation and subsequent mixing yield greater mucus clump disruption, facilitating CFU plating.

HBE cultures used in these infections were differentiated with and without PYO (0, 1, or 5 µM, as labeled in the figure) for 45-50 days. To enable a more accurate comparison, HBE cultures from the same batch were used for each experiment. For imaging experiments, HBE cultures were incubated overnight with CellMask Deep Red dye (Life Technologies, 5 µg/mL) to label the plasma membrane. The dye was added to the P-ALI medium on the basal side. The P-ALI containing the CellMask dye was then replaced with fresh P-ALI containing propidium iodide (PI, 2.5 µg/mL), which was used during infection to visualize tissue damage. No washes were performed from the apical side at any point during HBE differentiation or prior to infection.

## Data processing and plotting

With the exception of analyses related to scRNA-seq, data processing was primarily carried out in Python (version 3.13.5)(*85*), using the Pandas (version 2.2.3) (*86*), NumPy (version 1.26.4) (*87*), and SciPy (version 1.15.3) (*88*) libraries. Details of image analysis workflows are described in the corresponding section above. Microsoft Excel (version 16.105.1) was also used for selected analyses. The data were then plotted using Matplotlib (version 3.10.0) (*89*) and Seaborn (version 0.13.2) (*90*), with plot legends and their layout refined in Adobe Illustrator 2025 (version 29.5.1, Adobe), which was also used to create all schematic illustrations included in the manuscript.

## Statistical analyses

Except for analyses of the scRNA-seq data (see above), all statistical analyses shown in the figures were performed using Python. In these analyses, either Welch unpaired t-tests (two-sided) or one-way analysis of variance (ANOVA) with post hoc Tukey honestly significant difference (HSD) test for multiple comparisons were used, as specified in the figures’ legends.

