## Supplementary Material for "Pyocyanin disrupts airway regeneration to favor chronic *Pseudomonas aeruginosa* infection"

### Supplementary Material for Meirelles et al.

#### Supplementary Figures

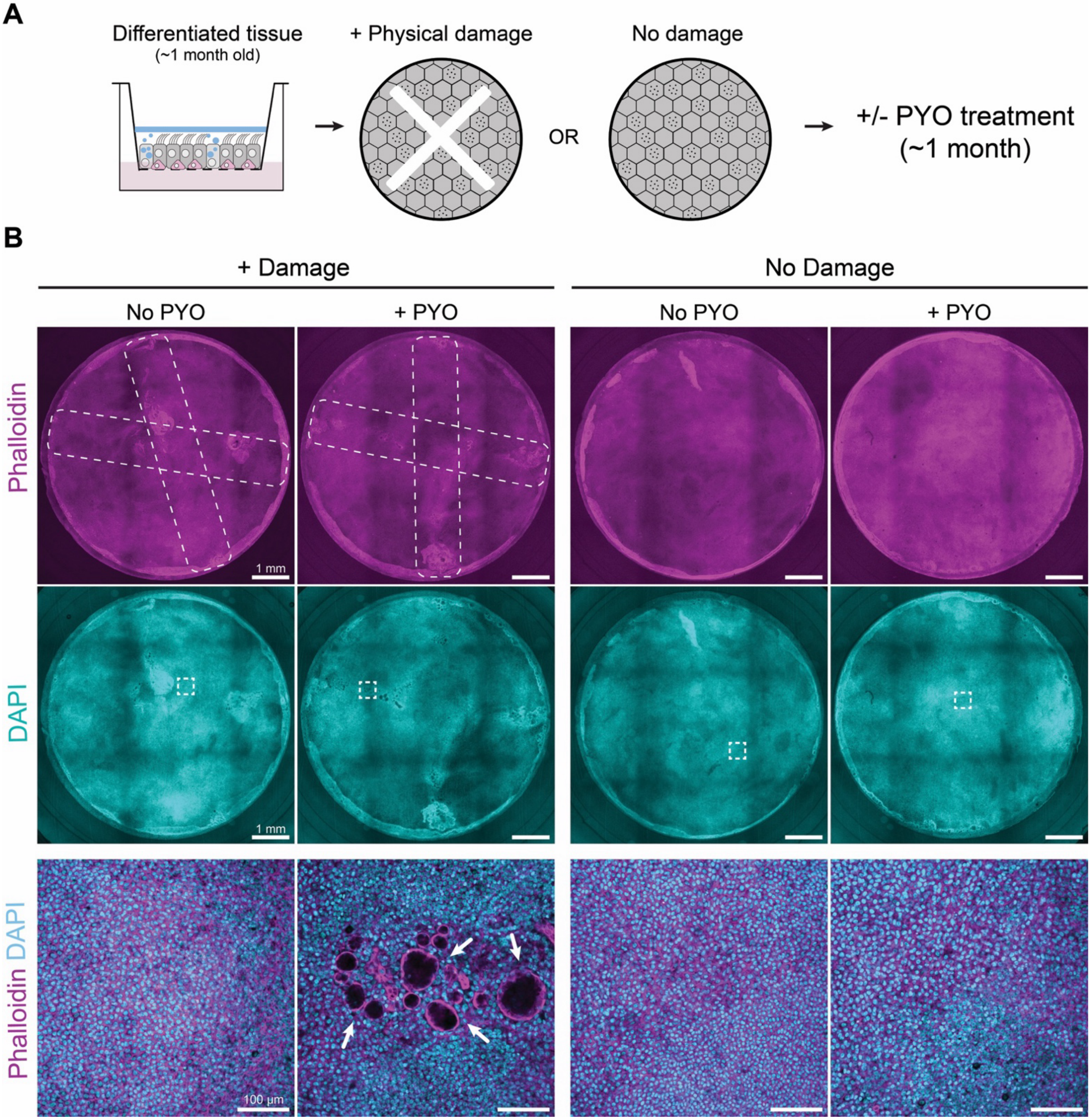


**Figure S1. The bacterial metabolite PYO disrupts airway regeneration. A.** Experimental design used to test the effect of PYO during regeneration of the airway epithelium upon physical damage. Briefly, fully differentiated HBE cultures on Transwells with or without physical damage were exposed to PYO at the basal side. **B.** Confocal microscopy of HBE cultures subjected to the different conditions tested after 29 days of exposure to PYO (5 µM). Rectangular dashed lines represent the approximate area that was damaged, and smaller squares highlight the approximate area visualized at higher magnification in the bottom images. Multiple cysts (white arrows) were often observed in the recovered area exposed to PYO. To improve the display of morphological features, brightness and contrast were adjusted independently for each image.

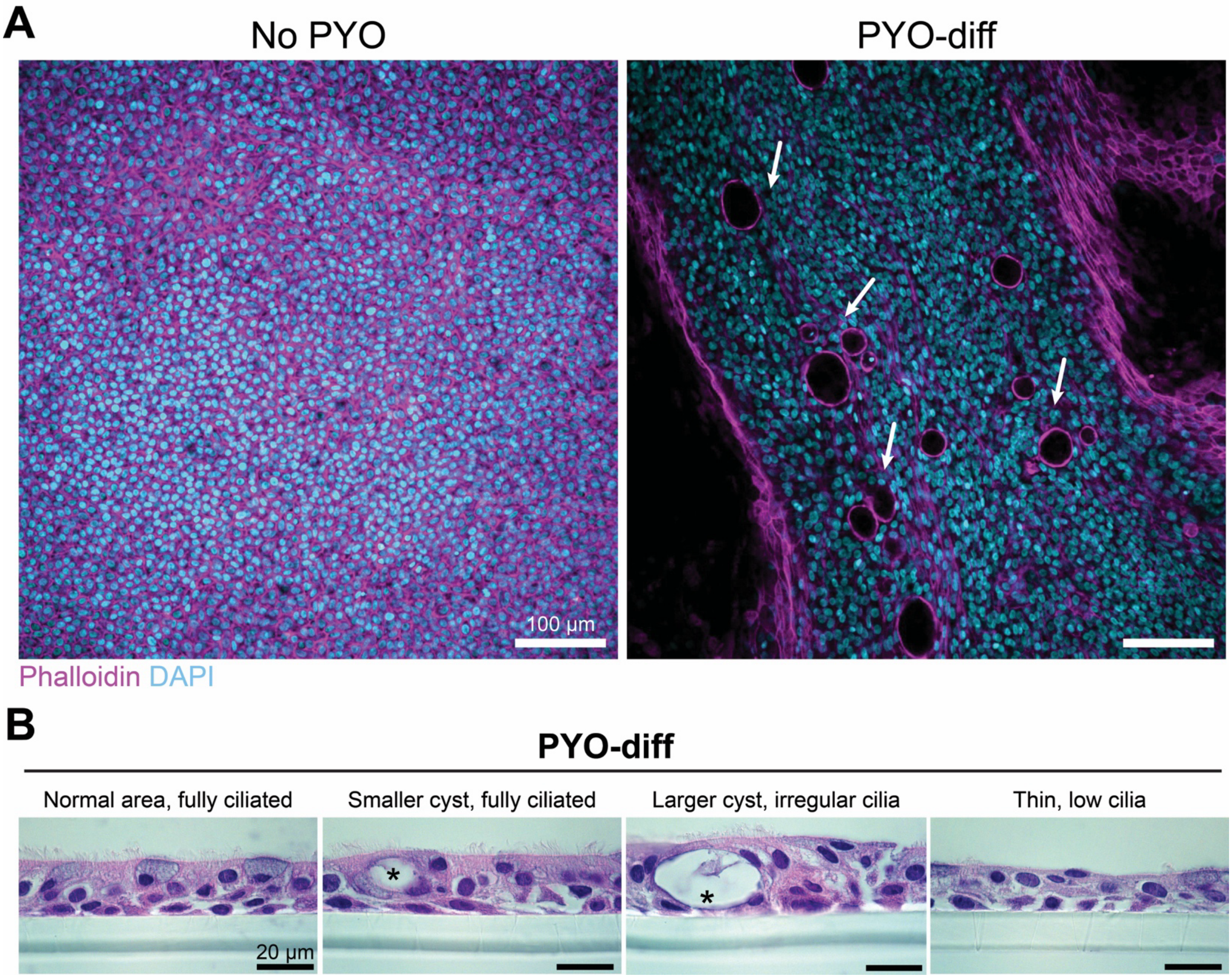


**Figure S2. Tissue morphology upon PYO exposure during differentiation. A.** Confocal microscopy of HBE cultures (generated from primary cells from a CF donor) after differentiation under PYO exposure (1 µM). As in cultures from a healthy donor, PYO induced tissue thickening and the emergence of cysts (white arrows). The image on the right shows a z-slice through the middle of the tissue thickening, located between two thinner areas (see Movie S12-13 for comparison). These z-stacks were used for the 3D reconstruction in Fig. 1E. **B.** Additional histology slides (Hematoxylin and eosin - H&E - stain) of tissues differentiated with PYO (5 µM). The asterisks mark cysts. PYO exposure during differentiation leads to tissue heterogeneity, and the four images represent distinct morphologies found in different areas of the tissue (the same image as in Fig. 1F is included for comparison).

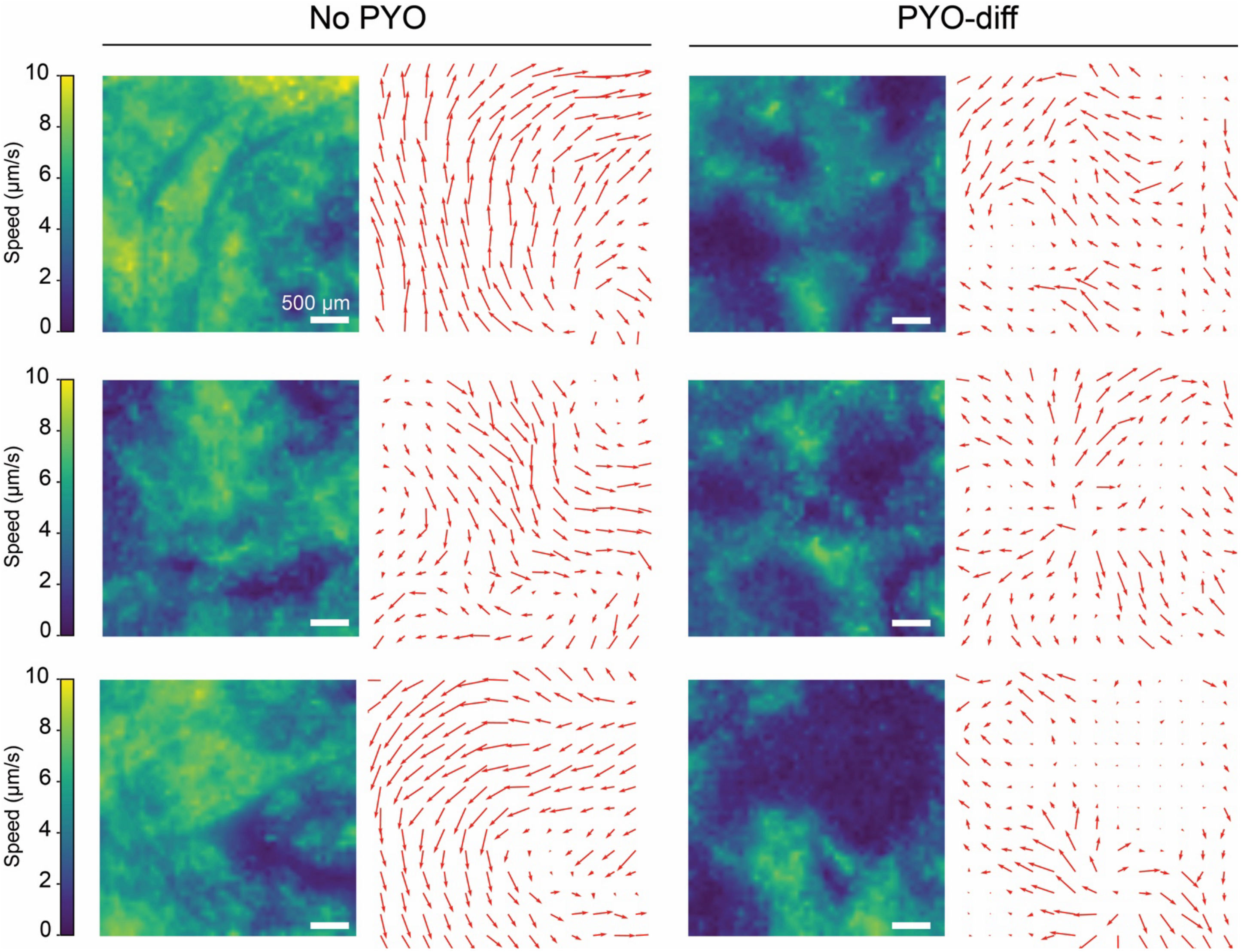


**Figure S3. PYO exposure during differentiation creates a tissue with heterogeneous mucociliary clearance.** Heatmaps of flow speed (left) or velocity vector fields (right) for all three replicates tested. Differentiation under PYO (5 µM) increases flow heterogeneity. To assess only the long-term effects of PYO (i.e., differences in tissue structure, not immediate physiological changes), the molecule was removed ~24 hours before the experiment.

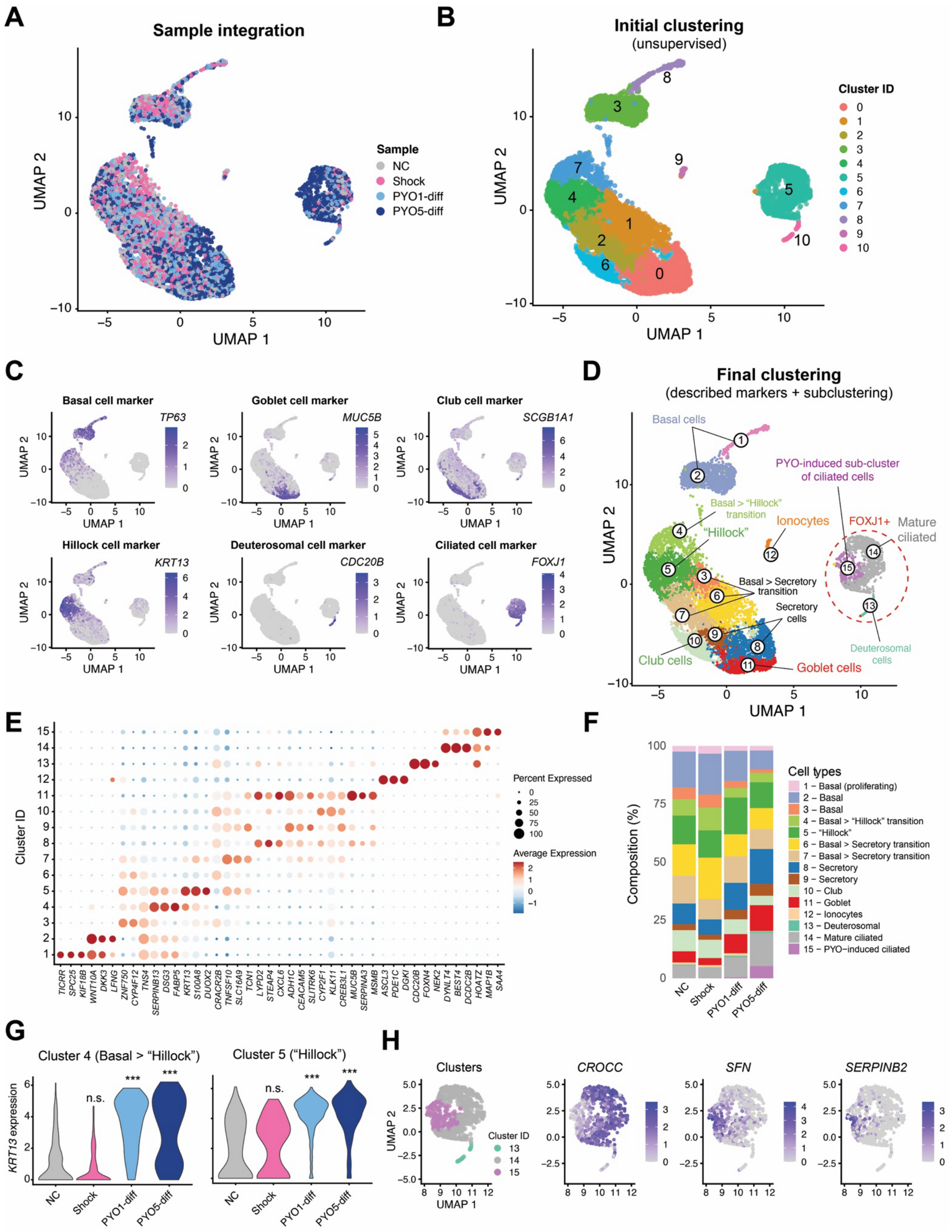


**Figure S4. Studying the effect of PYO on differentiation using single-cell RNA sequencing (scRNA-seq). A-D.** Analysis steps to identify cell clusters (10163 cells). After preprocessing and integration (A), 11 initial clusters (0-10) were identified using an unsupervised Louvain algorithm (B); next, we used expression of canonical markers for the expected cell types present in the airway epithelium to identify the cell type represented by each cluster (C); finally, based on the expression patterns of these genes, we performed final sub-clustering, resulting in the 15 clusters we report in our analyses (D). **E.** Top 3 expression-based markers from our dataset for each of the 15 identified clusters. For a full list of markers and their scores, see Table S9. Importantly, these markers were not necessarily used for annotation. For this purpose, we used literature-curated markers. **F.** Composition of cell-type populations within each sequenced sample. **G.** *KRT13* expression levels across the four conditions studied, considering only cells from clusters 4 (left) and 5 (right). The expression of cells from these two clusters combined is shown in Fig. 2F. **H.** Expression of three key genes related to cilia assembly (CROCC) and squamous and hillock identity (SFN and SERPINB2, respectively) was identified during differential gene expression analysis between cluster 15 (PYO-induced) and cluster 14 (Mature ciliated) using cells from the PYO5-diff sample. Statistics in **G**, Wilcoxon Rank Sum test after Benjamini-Hochberg correction for multiple comparisons, with asterisks showing significant differences relative to NC (***$p < 0.001$; n.s., $p > 0.05$)

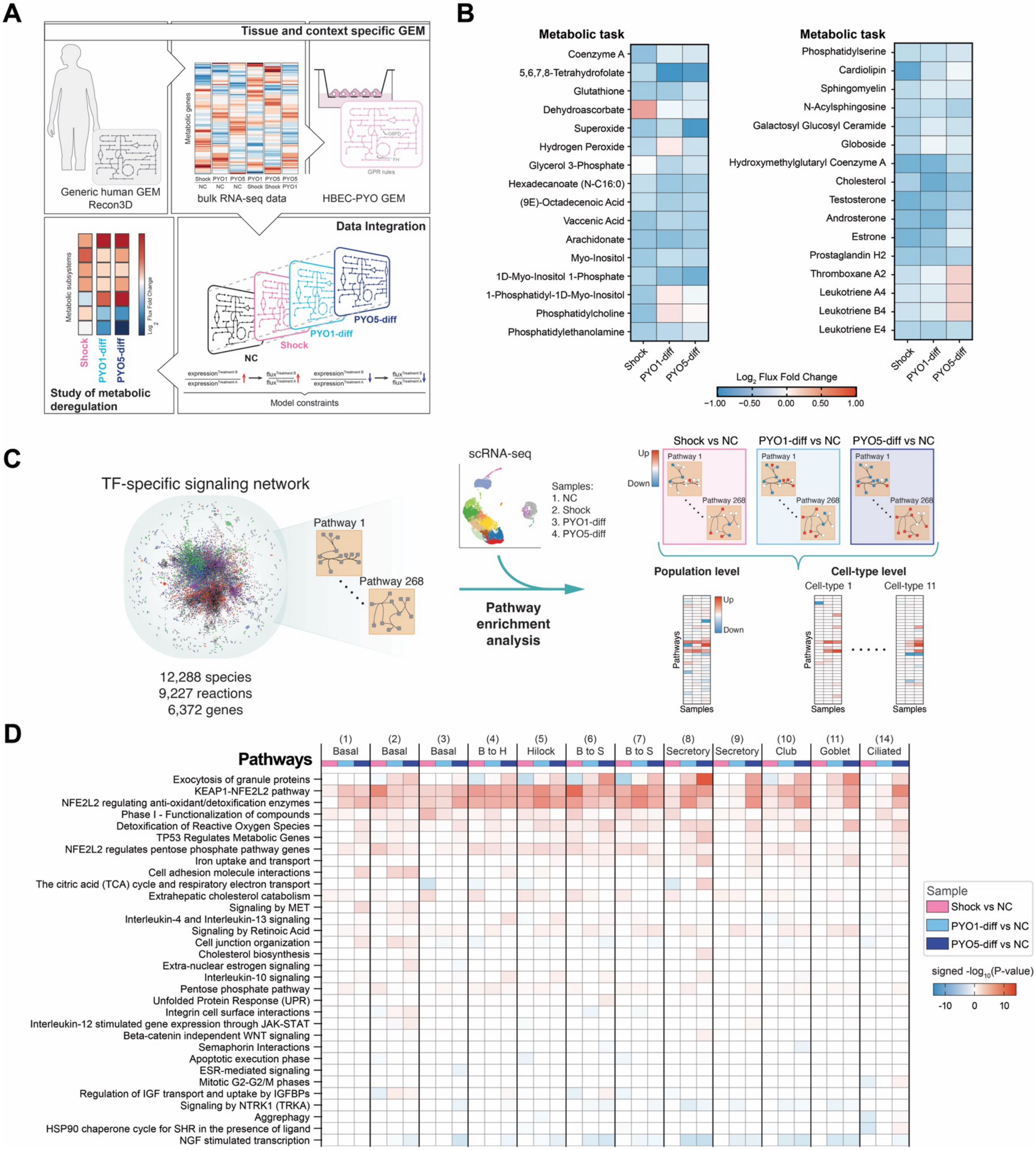


**Figure S5. PYO-diff effects on metabolism and signaling. A.** Schematic overview of the metabolic modeling workflow. The human genome-scale metabolic network Recon3D was reduced using the redHUMAN framework to generate a HBEC-specific, PYO-contextualized metabolic model (redHBECPYO), focusing on metabolic subsystems enriched in control cells and/or deregulated upon PYO exposure. Pseudo-bulk transcriptomic data were integrated using an extended version of the REMI method, enabling the simultaneous incorporation of pairwise gene expression-derived flux ratios across multiple conditions. This approach identified flux distributions consistent with network constraints and gene-expression changes, thereby enabling the prediction of PYO-induced metabolic deregulation. The constrained solution space was sampled to obtain representative reaction fluxes for downstream analyses. **B.** Fold change of metabolic task activities in PYO-treated conditions relative to the negative control. Task activities were computed from representative reaction fluxes within task-specific minimal functional networks, considering only core reactions shared across all alternative minimal networks for each task. **C.** Workflow for signaling analysis. We mapped differential expression from single-cell RNA sequencing onto a mechanistic signaling network centered on transcription factors that regulate metabolic genes and organized into 268 Reactome-annotated pathways. For each of the three treatments versus the negative control, we performed pathway enrichment analysis using an over-representation test applied to (i) population-level and (ii) cell-type-resolved differential expression results. This analysis identified the signaling programs most deregulated by each PYO treatment and linked these upstream changes to downstream metabolic reprogramming. **D.** Enriched signaling pathways at the cell-type level for each of the three treatments versus the negative control. Heatmap displays Benjamini-Hochberg adjusted $-\log_{10}$ ($p$-values) calculated via upper-tailed hypergeometric test, signed to indicate enrichment of upregulated (red) or downregulated (blue) genes.

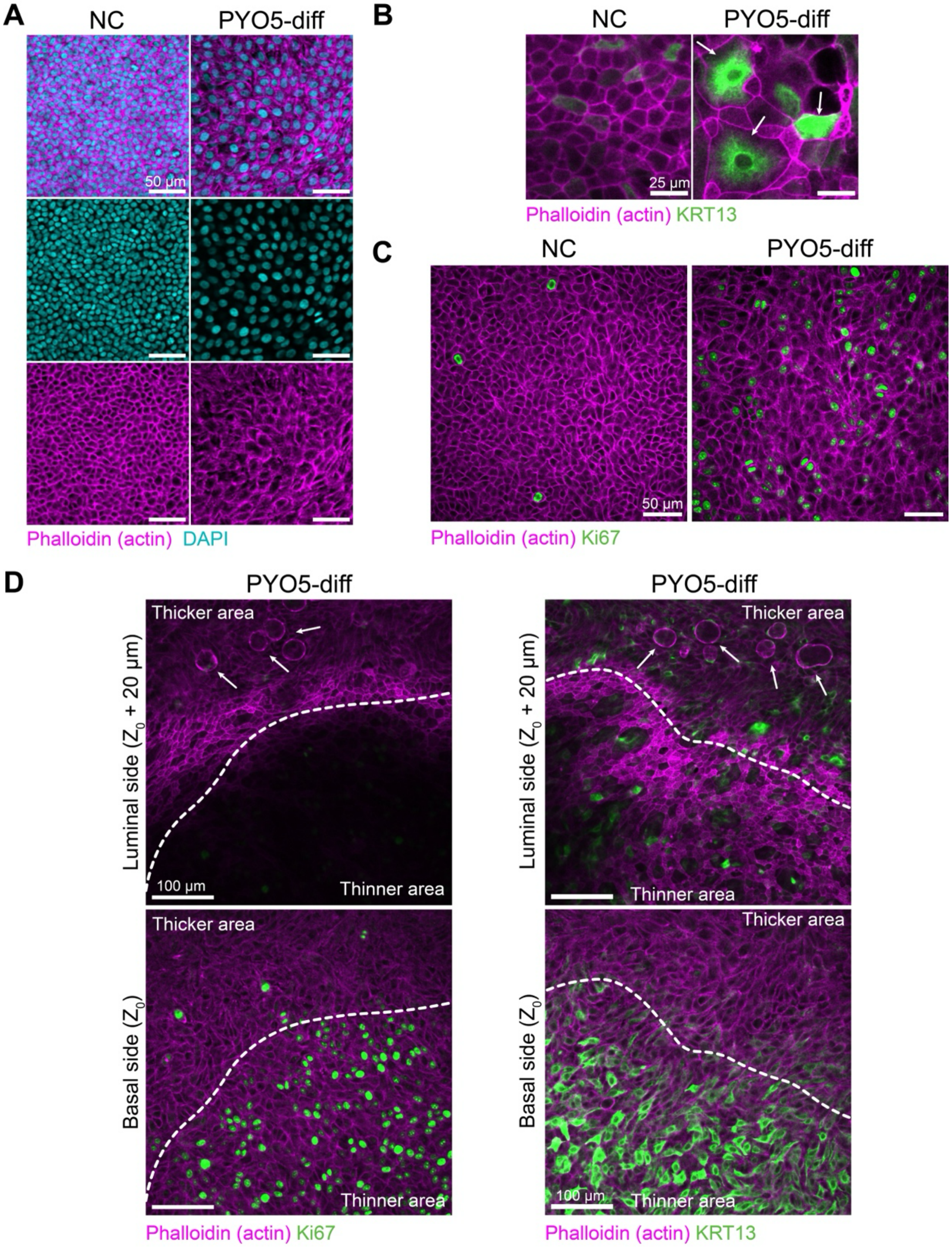


**Figure S6. Squamous and proliferative phenotypes induced by the PYO-diff treatment. A.** Representative images showing the basal side of airway epithelium in the NC or in a patch of PYO-induced disrupted cilia. These represent the basal side of areas shown in Fig. 3C. Note the squamous morphology in the PYO5-diff sample, with elongated cells occupying a larger area in the xy plane. **B.** Representative IF images of the luminal side of airway epithelium in the NC or in a patch of PYO-induced disrupted cilia, which often contains multiple enlarged flat cells with high KRT13 expression (see also Fig. 3K). **C.** Representative IF images of the basal side of airway epithelium in the NC or in a patch of PYO-induced disrupted cilia, which often contains multiple $Ki67^+$ cells on the basal side, indicating proliferation (see Movies S14 and S15 for the full z-stacks, respectively). **D.** Higher magnification showing the mirrored patterns of the location of proliferative ($Ki67^+$; left) and $KRT13^+$ cells (right) adjacent to thicker tissue in two distinct PYO5-diff samples (see Movies S9 and S16 for the full z-stacks, respectively). Arrows point to cysts found in the thicker part of the tissue.

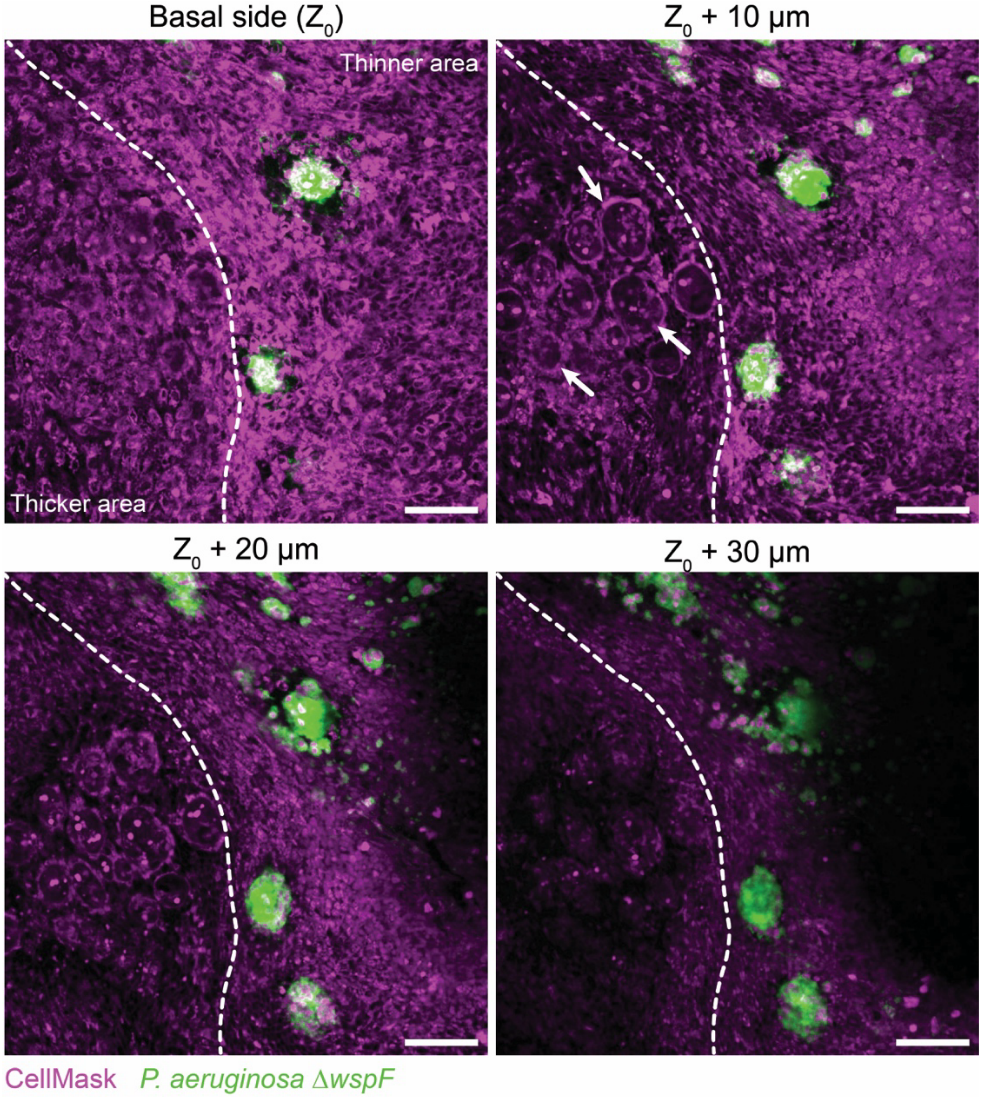


**Figure S7.** ***P. aeruginosa*** **hyperbiofilm mutant growth in PYO5-diff tissues.** Representative area showing Δ*wspF* growth at the mucosal surface, from the basal side ($Z_0$) to the luminal side ($Z_0$ + 30 µm). Growth is often localized in thinner regions adjacent to thicker areas of the tissue. Arrows in the CellMask channel point to cysts found in the thicker part of the tissue (see Movie S17 for the full z-stack).

#### Supplementary Tables

**Table S1. Results of differential gene expression (DGE) analysis using the pseudo-bulk approach**. In this case, DGE was performed for the entire population (i.e., without considering cell clusters) across the 3 conditions (Shock, PYO1-diff, PYO5-diff) relative to the control (NC).

**Table S2. Results of metabolic modeling.** This table presents the gene deregulation, alternative minimal networks generated for each defined metabolic task, and the representative flux values for all reactions in the redHBECPYO model.

**Table S3. Results of cluster-specific differential gene expression (DGE) analysis.** In this case, DGE was performed in each cluster across the 3 conditions (Shock, PYO1-diff, PYO5-diff) relative to the control (NC).

**Table S4. Results of signaling enrichment analysis.** This table presents the results of signaling pathway enrichment at the population level (Tabs a and b) and at the cell-type level (Tabs c and d). Shown values are FDR-adjusted signed $-\log_{10}$(p-values) for each treatment vs the negative control.

**Table S5. Results of Tn-seq analysis comparing distinct samples with the inoculum.** This table presents the output of TRANSIT for fitness ($\text{Log}_2\text{FC}$) and the adjusted p-value for each gene in the *P. aeruginosa* genome. Results are shown for the library grown at the mucosal surface (NC, PYO1-diff, PYO5-diff) and in rich media (P-ALI, LB) relative to the liquid culture inoculum reference.

**Table S6. Results of Tn-seq analysis comparing PYO1-diff or PYO5-diff treatments with NC.** This table presents the output of TRANSIT for fitness ($\text{Log}_2\text{FC}$) and the adjusted p-value for each gene in the *P. aeruginosa* genome. Results are shown for the library grown at the mucosal surface of PYO-exposed HBE (PYO1-diff and PYO5-diff) versus the mucosal surface of untreated HBEs (Negative untreated control – NC).

**Table S7. Functional annotation of Tn-seq data using the Database for Annotation, Visualization and Integrated Discovery (DAVID).** This table presents the results of enrichment analysis using the DAVID web tool for KEGG Pathways.

**Table S8. Results of one-way ANOVA assessing variability of insertion counts across conditions.** This table presents the results of one-way analysis of variance (ANOVA) testing genes that exhibit statistically significant variability in insertion counts across multiple conditions (NC, PYO1-diff, PYO5-diff, P-ALI, and LB).

**Table S9. List of markers for each cluster.** This table lists all markers assigned to each cluster using the Seurat FindAllMarkers function. Each tab corresponds to a specific cluster. Fig. S4e shows the top 3 markers for each cluster based on expression level.

**Table S10. Strains and primers used in this study.**

#### Supplementary Movies

**Movie S1.** A representative *z*-stack of a damaged area of HBE after regeneration in the absence of PYO (control). The movie progresses from the basal to the apical surface of the epithelium at 1 µm per step. Phalloidin (actin) signal is shown in purple, while DAPI (DNA) signal is shown in light blue. A *z*-slice derived from this data is shown in Fig. 1B-left.

**Movie S2.** A representative *z*-stack of a damaged area of HBE after regeneration in the presence of 5 µM PYO. The movie focuses on an area of increased thickness and progresses from the basal to the apical surface of the epithelium at 1 µm per step. Phalloidin (actin) signal is shown in purple, while DAPI (DNA) signal is shown in light blue. A *z*-slice derived from this data is shown in Fig. 1B-right.

**Movie S3.** A representative *z*-stack of HBE morphology after differentiation in the absence of PYO (NC). The movie progresses from the basal to the apical surface of the epithelium at 1 µm per step. Phalloidin (actin) signal is shown in purple, while DAPI (DNA) signal is shown in light blue. A *z*-slice derived from this data is shown in Fig. 1D-left.

**Movie S4.** A representative *z*-stack of HBE morphology after differentiation in the presence of 5 µM PYO (PYO-diff). The movie focuses on an area of increased thickness and progresses from the basal to the apical surface of the epithelium at 1 µm per step. Phalloidin (actin) signal is shown in purple, while DAPI (DNA) signal is shown in light blue. A *z*-slice derived from this data is shown in Fig. 1D-right.

**Movie S5.** A representative movie of the mucociliary flow generated by HBE differentiated in the absence of PYO (control). The movie spans 30 seconds, and the green signal comes from 2 µm-wide fluorescent beads used in the assay. This data was used to generate Fig. 1H-left.

**Movie S6.** A representative movie of the mucociliary flow generated by HBE differentiated in the presence of 5 µM PYO (PYO-diff). The movie spans 30 seconds, and the green signal comes from 2 µm-wide fluorescent beads used in the assay. This data was used to generate Fig. 1H-right.

**Movie S7.** A representative *z*-stack of KRT13 and acetylated-α-tubulin signal in HBE after differentiation in the presence of 5 µM PYO (PYO5-diff). The movie focuses on a squamous area with disrupted cilia distribution and progresses from the basal to the apical surface of the epithelium at 1 µm per step. Phalloidin (actin) signal is shown in purple, KRT13 in green, and acetylated-α-tubulin in orange. *z*-slices derived from this data are shown in Fig. 3K-right.

**Movie S8.** A representative *z*-stack of KRT13 and acetylated-α-tubulin signal in HBE after differentiation in the absence of PYO (NC). The movie progresses from the basal to the apical surface of the epithelium at 1 µm per step. Phalloidin (actin) signal is shown in purple, KRT13 in green, and acetylated-α-tubulin in orange. *z*-slices derived from this data are shown in Fig. 3K-left.

**Movie S9.** A representative *z*-stack of Ki-67 signal in HBE after differentiation in the presence of 5 µM PYO (PYO5-diff). The movie focuses on a squamous region adjacent to a region of increased thickness and progresses from the basal to the apical surface of the epithelium at 1 µm per step. Phalloidin (actin) signal is shown in purple, while Ki-67 signal is shown in green. *z*-slices derived from this data are shown in Fig. 4C and Fig. S6D-left.

**Movie S10.** A representative *z*-stack of a zoomed-in area showing Δ*wspF* growth at the mucosal surface of an HBE after differentiation in the absence of PYO (NC). The movie progresses from the basal to the apical surface epithelium at 1 µm per step. CellMask Deep Red (plasma membrane of epithelial cells) signal is shown in purple, while Δ*wspF P. aeruginosa* signal is shown in green. A *z*-slice derived from this data is known as Fig. 6B-top.

**Movie S11.** A representative *z*-stack of a zoomed-in area showing Δ*wspF* growth at the mucosal surface of an HBE after differentiation in the presence of 5 µM PYO (PYO5-diff). The movie progresses from the basal to the apical surface epithelium at 1 µm per step. CellMask Deep Red (plasma membrane of epithelial cells) signal is shown in purple, while Δ*wspF P. aeruginosa* signal is shown in green. A *z*-slice derived from this data is shown in Fig. 6B-bottom.

**Movie S12.** A representative *z*-stack of CF HBE morphology after differentiation in the absence of PYO (NC). The movie progresses from the basal to the apical surface epithelium at 3 µm per step. Phalloidin (actin) signal is shown in purple, while DAPI (DNA) signal is shown in light blue. A *z*-slice derived from this data is shown in Fig. S2A-left.

**Movie S13.** A representative *z*-stack of CF HBE morphology after differentiation in the presence of 1 µM PYO (PYO-diff). The movie focuses on an area of increased thickness and progresses from the basal to the apical surface of the epithelium at 3 µm per step. Phalloidin (actin) signal is shown in purple, while DAPI (DNA) signal is shown in light blue. A *z*-slice derived from this data is shown in Fig. S2A-right.

**Movie S14.** A representative *z*-stack of Ki67 signal in HBE after differentiation in the absence of PYO (NC). The movie progresses from the basal to the apical surface epithelium at 1 µm per step. Phalloidin (actin) signal is shown in purple, while Ki67 signal is shown in green. A *z*-slice derived from this data is shown in Fig. S6C-left.

**Movie S15.** A representative *z*-stack of Ki67 signal in HBE after differentiation in the presence of 5 µM PYO (PYO5-diff). This movie focuses on a patch with disrupted cilia and progresses from the basal to the apical surface epithelium at 1 µm per step. Phalloidin (actin) signal is shown in purple, while Ki67 signal is shown in green. A *z*-slice derived from this data is shown in Fig. S6C-right.

**Movie S16.** A representative *z*-stack of KRT13 signal in HBE after differentiation in the presence of 5 µM PYO (PYO5-diff). As in Movie S9, this movie focuses on a squamous region adjacent to a region of increased thickness and progresses from the basal to the apical surface of the epithelium at 1 µm per step. Phalloidin (actin) signal is shown in purple, while KRT13 signal is shown in green. *z*-slices derived from this data are shown in Fig. S6D-right.

**Movie S17.** A representative *z*-stack of a zoomed-in area showing Δ*wspF* growth at the mucosal surface of an HBE after differentiation in the presence of 5 μM PYO (PYO5-diff). The movie focuses on an area with abundant bacteria in squamous regions adjacent to thicker tissue containing cysts, and progresses from the basal to the apical surface of the epithelium at 1 μm per step. CellMask Deep Red (plasma membrane of epithelial cells) signal is shown in purple, while Δ*wspF P. aeruginosa* signal is shown in green. *z*-slices derived from this data are shown in Fig. S7.
